# A Generative Foundation Model for Antibody Design

**DOI:** 10.1101/2025.09.12.675771

**Authors:** Rubo Wang, Fandi Wu, Jiale Shi, Yidong Song, Yu Kong, Jian Ma, Bing He, Qihong Yan, Tianlei Ying, Peilin Zhao, Xingyu Gao, Jianhua Yao

## Abstract

Antibodies are indispensable components of the immune system, yet the design of high-affinity antibodies remains a time-consuming and experimentally intensive process. To address this challenge, we present IgGM, a novel generative foundation model designed to accelerate high-affinity antibody engineering. IgGM learns the complex relationships underlying the binding interactions between antigens and antibodies, as well as the mapping between antibody sequences and structures. By conditioning on different inputs, IgGM supports a wide range of antibody design tasks, including complex structure prediction, inverse design, affinity maturation, framework optimization, humanization, and *de novo* antibody design. It is compatible with both conventional antibodies and nanobodies, and allows user-defined CDR loop lengths for flexible design. To prioritize candidates, we introduce a frequency-based computational screening strategy that enhances design efficiency. Extensive evaluation through both *in silico* benchmarks and *in vitro* experiments across diverse antigens such as PD-L1, Protein A, TNF-***α***, IL-33, SARS-CoV-2 RBD and its variants demonstrates that IgGM consistently generates antibodies or nanobodies with high measured affinity. These results underscore IgGM’s versatility and effectiveness as a powerful tool for next-generation antibody discovery and optimization.

## 1 Introduction

Antibodies, also known as immunoglobulins (Ig), are Y-shaped proteins secreted by B lymphocytes, primarily present in blood and lymphatic fluid [1, 2]. Structurally, an antibody consists of two heavy chains and two light chains, each containing a variable domain (VH for heavy chains, VL for light chains) and a constant domain (CH for heavy chains, CL for light chains). The variable regions are composed of three complementarity-determining regions (CDRs) and four framework regions (FRs). CDRs are critical for antigen binding and define antibody specificity, while FRs provide structural stability to the variable regions (VR) with relatively low variability. Antibodies play a pivotal role in the immune system: they recognize and bind to specific foreign entities (*e*.*g*., bacteria, viruses, fungi, and parasites) and mark these invaders for elimination [3, 4]. Beyond immunity, their applications extend to medicine, scientific research, and biotechnology, including disease treatment, personalized medicine, vaccine development, and novel drug discovery[5–7]. The core mechanism underlying their function lies in antibody-antigen interaction, and antibody design encompasses diverse scenarios such as *de novo* design, affinity maturation, humanization, and framework region engineering.

Traditional antibody production methods, despite their foundational role, face significant challenges. As shown in Fig. 1a, these include long production cycles [8], batch-to-batch variability [9], and the need for humanization to reduce immunogenicity [10]. Such limitations hinder their widespread application and restrict therapeutic efficacy. The advancement of deep learning has driven substantial progress in key tasks of antibody development. In the field of structure prediction, tools based on multiple sequence alignment (MSA), such as AlphaFold3 [11] and AlphaFold-Multimer [12], and pre-trained language models, such as IgFold [13], ABlooper [14], and tFold [15]), have achieved rapid and highly accurate prediction of antibody and antibody-antigen complex structures. These tools enable precise prediction of antibody structures and antibody-antigen complexes, with exceptional performance in conformational modeling of CDRs, particularly the H3 loop. Complementing structure prediction is its inverse problem: inverse design. Inverse design methods [16–19] focus on redesigning CDR sequences with targeted functions, such as generating antibodies with higher binding affinity or bypassing patent restrictions. Notably, these methods achieve such design goals by leveraging known antibody structures as a starting point, rather than*de novo*generation from scratch.

**Fig. 1:**
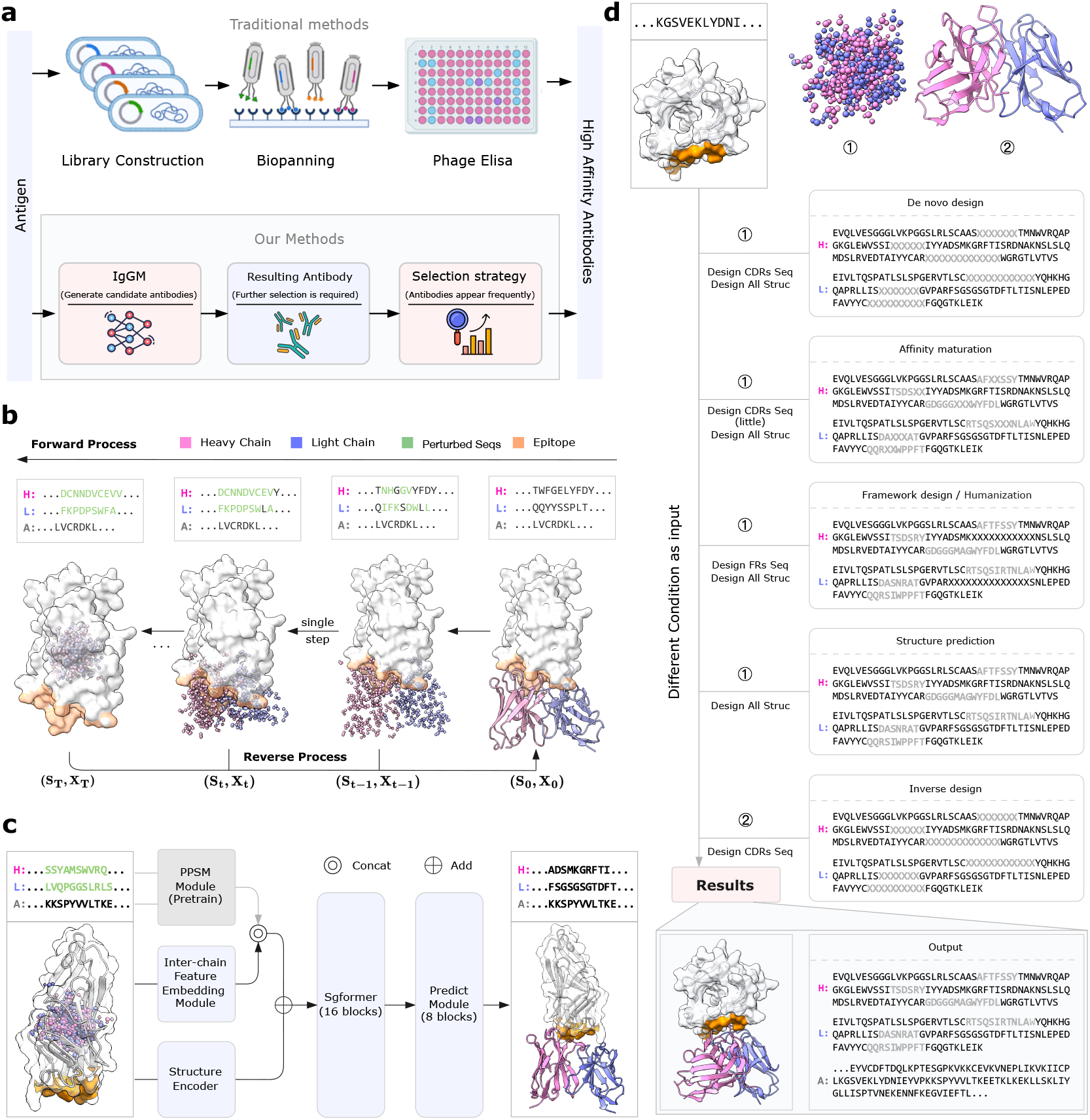
IgGM is a versatile generative model capable of designing different functional regions. **a**, Traditional antibody engineering methods involve complex processes, such as combinatorial library construction, screening, and identification. Our method uses IgGM to generate candidate antibodies against specific antigens. Subsequently, the frequency of antibody occurrence is used as a screening strategy to obtain antibodies with high affinity. **b**, The forward process and reverse process for co-design of antibody sequences and structures: The figure illustrates an example trajectory in the design of antibodies, incorporating specific antigen and antibody. It depicts the input data (***s***_*T*_, ***x***_*T*_) to the model, along with the corresponding predicted outcomes (***s***_0_, ***x***_0_). The consistency model is capable of generating antibody sequences and structures across diverse noise levels, demonstrating the gradual refinement from noisy inputs to a well-defined antibody structure. Gray represents the sequence region of CDR, and ‘X’ indicates the region that needs to be designed. The structure of different forms of antibody represents the initial structural input, which varies under different conditions. **c**, IgGM model framework diagram: Before being input into the model, the antibody sequence and structure are pre-noised. The model includes a pre-trained PPSM module, a Inter-chain Feature Embedding Module, a Structure Encoder, a Feature Embedding Module, 16 layers of Sgformer, and an 8 layers of Prediction Module. The model is able to predict and recover the structure and sequence. **d**, IgGM is a multifunctional antibody generation model that can be applied in various antibody design scenarios taking antigen/epitope information as inputs and providing different conditional input. It can be applied in *de novo* antibody design, affinity maturation, framework region design, structure prediction, as well as inverse design. Different design scenarios correspond to different input information.

Beyond structure prediction and inverse design, deep learning has also made significant strides in other critical areas of antibody development. For affinity maturation, the self-supervised model GearBind [20] predicts the binding free energy change (ΔΔG_bind_) by extracting the geometric representations of wild-type and mutant antibodies, successfully optimizing the affinity of the anti-SARS-CoV-2 antibody CR3022. In antibody design, models such as RefineGNN [21] adopt a sequence and structure co-design strategy, integrating graph neural networks to iteratively refine global structures and sequences. Meanwhile, RFantibody [22] generates an antibody backbone with shape complementarity driven by the target epitope, achieving “atomic-level accuracy *de novo* design” and overcoming traditional template dependence.

A critical limitation persists across current methodologies: no single framework comprehensively addresses the diverse spectrum of antibody design tasks while maintaining independence from pre-existing antibody structures. While numerous sequence-structure co-design methods [23–26] have validated AI-driven antibody design feasibility, most rely on known experimental structures of antibody-antigen complexes or modifications of pre-existing functional antibodies. When designing novel antibodies against specific antigen targets without prior structural knowledge, these approaches become inapplicable. Furthermore, existing models typically specialize in single tasks (*e*.*g*., structure prediction *or* affinity maturation) rather than providing a unified solution across multiple design scenarios. This fragmentation of capabilities creates workflow inefficiencies and limits the practical utility of computational antibody design in real-world therapeutic development.

To address these unmet needs, we propose IgGM, a novel generative model designed to tackle diverse antibody design tasks within a single unified framework. As illustrated in Fig. 1b, c, IgGM employs a multi-stage architecture: (1) leveraging a pre-trained protein language model (PPSM) to extract evolutionary sequence features for antibodies; (2) utilizing a feature encoder (Sgformer) to model antigen and antibody interactions; and (3) employing a prediction module to output both antibody sequences and structures. IgGM explicitly leverages the interplay between sequence and structure, enabling accurate design even from partial inputs—a capability critical for practical applications. Unlike previous approaches that require complete structural templates, IgGM can generate functional antibodies starting solely from antigen specifications. Specifically, it models antibody and antigen interactions, docks designed antibodies onto target epitopes, and generates sequences and structures of key regions (*e*.*g*., CDRs). Crucially, as summarized in Fig. 1d, by incorporating antigen/epitope information and supporting diverse conditional inputs, IgGM achieves exceptional versatility, covering key design scenarios including *de novo* design, affinity maturation, framework region engineering, humanization, and structure prediction within a single model. Furthermore, it supports both conventional antibodies and nanobodies, with user-specifiable CDR loop lengths. Experimental validation through both *in silico* analysis and wet-lab experiments across these tasks confirms that IgGM consistently generates antibodies exhibiting high binding affinity to target antigens, highlighting its broad applicability and potential for accelerating antibody discovery.

## 2 Results

### 2.1 IgGM is an accurate antigen-antibody complex structure prediction and an antibody inverse design tool

Antigen-antibody complex structure prediction and antibody inverse design represent two complementary capabilities of our framework that form a closed loop in antibody engineering. Complex structure prediction aims to generate the antigen-antibody complex structure using antigen/epitope information and the antibody sequence as a conditional input. Accurate prediction of this complex structure is fundamental for subsequent design tasks. Conversely, inverse design leverages structural information to generate optimal sequences, completing the design cycle. These capabilities jointly contribute to a comprehensive antibody engineering approach that closely links the relationships among sequence, structure, and function.

To execute complex structure prediction, IgGM utilizes the antigen/epitope data along with the full antibody sequence as input. Based on this input, the model directly predicts the 3D structure of the antigen-antibody complex as shown in Fig. A1a. To benchmark IgGM’s performance in complex structure prediction, we adopt the same evaluation metrics used by tFold-Ag [15]. Specifically, we used the RMSD of CDR3 and TM-Score [27] to evaluate the antibodies, and used DockQ [28] and the success rate (where DockQ *>* 0.23) to evaluate the antigen-antibody complexes.

We evaluated the structural prediction performance on the SAb23H2 test set, which is derived from the SAbDab database [29]. For detailed information, please refer to Section 4.1. We compared four methods: the antibody structure prediction methods IgFold [13] and tFold-Ag [15], as well as the latest protein structure prediction method AlphaFold 3 [11], alongside the antibody design method dyMEAN [30]. Since IgFold only supports the prediction of antibody structures, we utilized HDock [31] to dock the predicted antibody structures with the original antigen. For tFold-Ag and AlphaFold 3, which can directly predict the structure of antigen-antibody complexes, we used the antigen-antibody sequences as input. In the case of dyMEAN and IgGM, we predicted the structure of the antigen-antibody complex given the antigen structure, using the antibody sequence as input.

As shown in Fig. 2a and Fig. A1b, IgGM achieved lower RMSD values (2.14Å vs 2.25Å) and higher lDDT scores (0.89 vs 0.88) compared to dyMEAN, demonstrating superior antibody structure prediction despite not using template-based initialization. Although there is still a gap compared to specialized structure prediction methods, the results are overall quite close, indicating that our method has learned the distribution of antibody structures. At the same time, we evaluated the RMSD of CDR H3. As shown in Table B3, IgGM can also achieve relatively good prediction results. In terms of docking performance, as shown in Fig. 2b and Fig. A1c, our method surpassed both structure prediction methods and antibody design methods on DockQ, demonstrating that IgGM can achieve high docking performance. Additionally, it showed improved accuracy on iRMS and LRMS, with a success rate of 0.4667, significantly higher than dyMEAN’s 0.067. This indicates that IgGM is capable of capturing the interactions between antigens and antibodies effectively. When using the structures predicted by AlphaFold 3 as the initial input for IgGM instead of randomly initialized structures, IgGM demonstrated improved performance across all metrics, particularly in docking-related indicators. Specifically, utilizing AlphaFold 3 structures as initial inputs increased the success rate by 20 percentage points (from 0.4667 to 0.6667) compared to using randomly initialized structures. As shown in Fig. 2c, IgGM can accurately predict the structures of antigen-antibody complexes and nanobody-antigen complexes. As shown in Fig. 2d, for the inaccurately predicted structures by AlphaFold3, IgGM is capable of making corrections to yield more suitable structures. For example, for an antigen-antibody complex (pdb id: 8e6k), the DockQ of the structure predicted by AlphaFold3 is only 0.007. When this initial structure is input into IgGM, the DockQ of the output structure increases to 0.603, suggesting that IgGM can function in correcting existing structures.

**Fig. 2:**
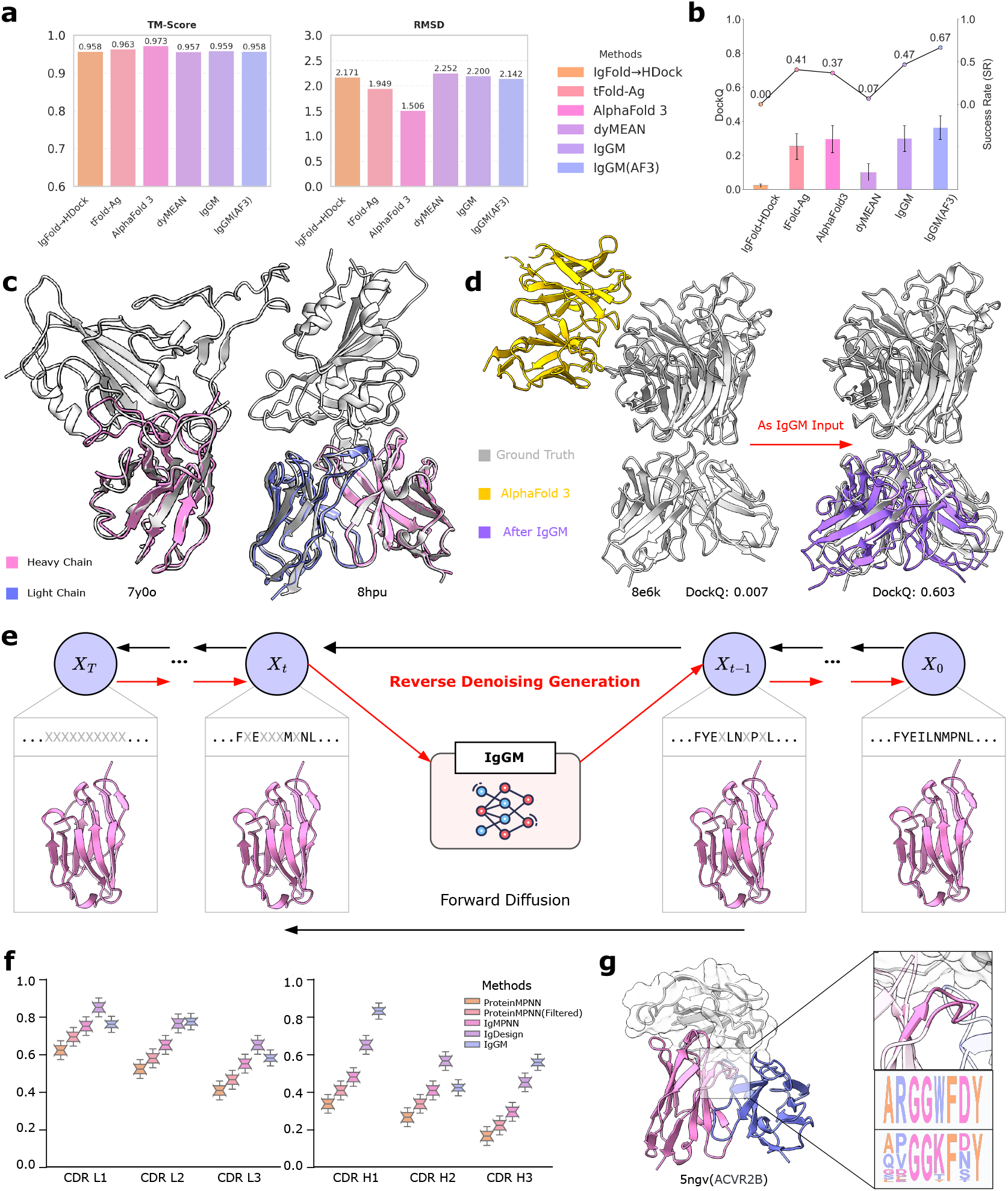
IgGM is an accurate complex structure prediction and an antibody inverse design tool. **a**, Performance of the structure prediction method on antibody structures. IgGM can outperform models other than AlphaFold 3. **b**, Performance of the structure prediction method on complex structures. IgGM can outperform other methods and achieve the highest prediction success rate. **c**, IgGM can predict accurate antigen-antibody complex structures and nanobody complex structures. Two examples are shown in the figure. **d**, IgGM can correct the errors predicted by AlphaFold 3. When the structure predicted by AlphaFold 3 deviates significantly from the epitope, IgGM can correct the output to a more accurate position.**e**. Flowchart of inverse design. The entire inverse design only involves changes in the sequence. **f**. Performance of different inverse design methods on the light-chain CDR and the heavy-chain CDR sequence recovery rate. **g**. The logo figure is about the original CDR3 sequence of ACVR2B (PDB ID: 5gnv) and the CDR3 sequence designed by IgGM.

This structural refinement capability naturally extends to the inverse design problem, where the goal shifts from structure prediction to sequence generation. While structure prediction takes sequence as input to generate structure, inverse design takes structure as input to generate optimal sequences, representing two sides of the same sequence-structure relationship that IgGM has learned to model comprehensively. Antibody inverse design is a computational method for predicting and designing antibody sequences based on the known structure of the antibody-antigen complex. Its core is to use a deep learning model to take the backbone structure of the antibody and the antigen-binding information as inputs, and generate amino acid sequences that can fold into the target structure. When the input of IgGM contains the initial complex structure, IgGM can effectively perform inverse folding to generate sequences that maintain the structural integrity of the target conformation. To optimize this capability, we implement a specialized fine-tuning procedure where the structure remains fixed while only the sequence is perturbed during training. As shown in Fig. 2e, only the sequence is noisy during input, and the structure remains unchanged throughout the process. We evaluated our model using the test set from IgDesign [18], ensuring experimental fairness by excluding related sequence clusters from the training data. The evaluation metric is AAR, which is used to evaluate the degree of similarity between the designed sequence and the original sequence. As shown in Fig. 2f, IgGM can outperform existing methods on CDR L2, CDR H1, and CDR H3, especially on the most important CDR H3. The performance metrics depicted in these figures highlight that IgGM achieves higher accuracy compared to ProteinMPNN, ProteinMPNN (Filtered), IgMPNN, and IgDesign across various CDR regions. Specifically, in Fig. 2f, IgGM demonstrates superior performance in CDR L2 with a median value close to 0.8, indicating its effectiveness in this region. Similarly, in Fig. 2g, IgGM shows significant advantages in CDR H1 and CDR H3, where it maintains a high median value above 0.6 and around 0.4, respectively. Although AAR only reflects the degree of similarity between designed antibodies and natural antibodies and cannot directly demonstrate the advantages of design performance, these results suggest that IgGM’s design and optimization strategies are particularly well-suited for capturing the complex structural and functional characteristics of these critical CDR regions, thereby enhancing the overall antibody design process.

### 2.2 IgGM can design novel and effective antibody framework regions

The design of the antibody framework region is a core step in antibody engineering. It involves multiple key tasks and aims to optimize the stability, developability, affinity of the antibody, and reduce its immunogenicity. Here we conduct verification for two tasks. One is to optimize the binding ability with protein A to enable large-scale production. The other is humanization, that is, to optimize the immunogenicity of the antibody so that mouse derived antibodies can be used in humans. Due to fundamental differences in the structural constraints and functional roles between framework regions (which provide structural stability) and CDR regions (which directly mediate antigen binding), the original IgGM model trained primarily on CDR-focused tasks cannot effectively design framework regions. To address this limitation, we implemented a targeted fine-tuning strategy specifically for framework region design. As shown in Fig. 3a, we curated a specialized dataset by extracting approximately 2,000 antigen-antibody complexes from the SAbDab structural database where framework regions directly contact the antigen (within 10 Å). During fine-tuning, we specifically perturb the framework regions in contact with the antigen by adding Gaussian noise to both sequence and structure, then train IgGM to restore these perturbed regions while maintaining overall structural integrity. This differs from the main training process in two critical aspects: (1) the noise application is targeted to framework regions rather than CDRs, and (2) the loss function emphasizes structural preservation of CDR regions during framework modifications. For these two tasks, we adopt distinct strategies for optimization. Subsequently, wet-lab experiments are carried out to verify the effectiveness of the framework region in relation to Protein A and the humanization of mouse antibodies.

**Fig. 3:**
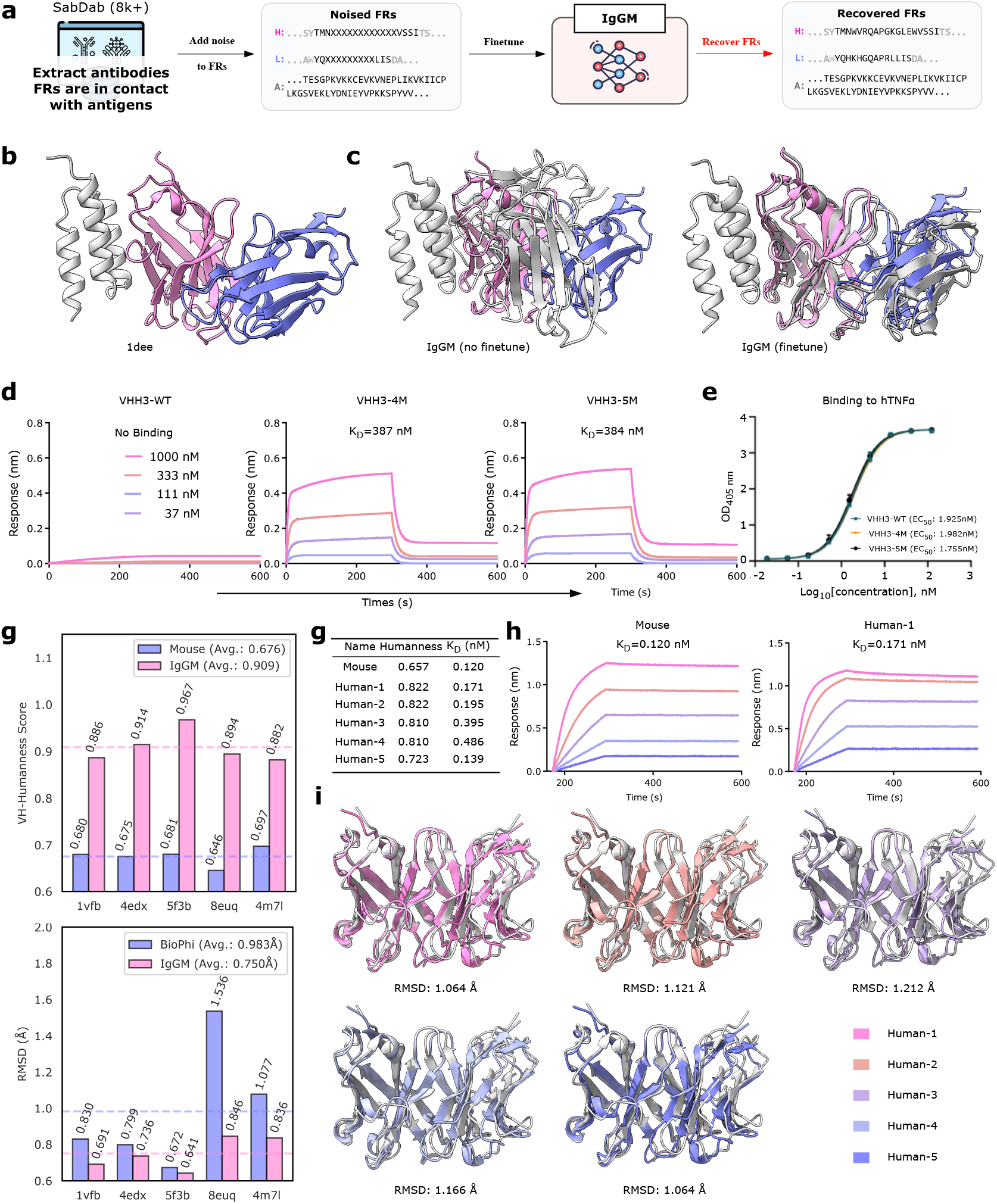
IgGM can be used for the design of the framework regions of antibodies. **a**, Before IgGM is applied to framework region design, it undergoes a fine-tuning process using antibodies whose framework regions are known to interact with antigens. This targeted dataset enables the model to better capture the structural and functional relevance of framework-antigen contacts. During training, controlled perturbations are introduced to both the framework region sequences and the overall antibody structure. **b**, The structural diagram of 1dee. 1dee contains a human IgM Fab that can interact with Protein A in terms of structure, and the binding region is within the framework region. **c**, The performance of IgGM before and after fine-tuning in the overall structure prediction of 1dee. The gray represents the original 1dee structure, and the color represents the predicted structure. Before fine-tuning, it will predict that the CDR region binds to Protein A, and fine-tuning can more accurately predict the binding position. DockQ increased from 0.05 to 0.88. **d**, The diagrams of the affinity curves for two effective variants were acquired through re-designing the binding framework region with IgGM and were measured by BLI. **e**,Binding curves of VHH3-4M, VHH3-5M, and VHH3-WT with human TNF alpha, aiming to test whether their original functions are retained. The EC50 values were reported. Among them, the EC50 of VHH3-WT was 1.925 nM, that of VHH3-4M was 1.982 nM, and that of VHH3-5M was 1.755 nM. The changes were not significant. **f**, Performance on multiple humanized test samples with structures. Above is the comparison of antibody humanization scores after IgGM optimization, and below is the comparison of RMSD between the IgGM optimized sequence and the BioPhi-generated sequence on the AlphaFold3 predicted structure. **g**, The results of the affinity curves for the antibody after IgGM humanization and were measured by BLI. **h**, Affinity curve graph of mouse derived antibody and Human-1. **i**, A comparison of the structural differences between the humanized antibody and the original antibody shows that they look very similar.

We first verified on antibodies that can achieve the combination of the framework region, and the target protein selected was Protein A. Protein A can bind to the framework regions of some antibodies, and at the same time, it can be effectively purified using commercial protein A resins, achieving higher purity and comparable yields to IMAC in a single step. The n501 is a fully human single domain antibody that targets the oncofetal antigen 5T4 [32]. The structure of the binding complex of n501 and Protein A is shown in the figure Fig. 3b (pdbid:1dee), and this binding affinity can reach the nanomolar level. We tested whether finetuning would have an effect on the n501 Protein A complex. As shown in Fig. 3c, the fine-tuned IgGM successfully predicts framework-region binding to Protein A, contrasting with the CDR-binding mode predicted by the unmodified model. As shown in Fig. A2a, VHH3 (PDB ID: 5M2M) is a single domain antibody that binds to TNF alpha [33]. Since its framework region cannot bind to Protein A—and no known single-domain antibody has been shown to do so—we selected this antibody as the basis for framework region design. Specifically, we redesigned the FR3 region of VHH3. Due to the high similarity among framework regions, the resulting antibodies exhibited only modest sequence variation. In total, we generated 1000 redesigned antibodies through iterative sampling. Antibodies with higher generation frequencies represent more probable solutions within the sequence-structure distribution learned by IgGM. We selected the top 10 antibodies with highest generation frequencies for wet-lab verification. As shown in Fig. 3d, BLI measurements confirmed that 2 of the 10 tested variants demonstrated specific binding to Protein A, with KD values of 387 nM and 384 nM respectively, while the remaining 8 variants showed no detectable binding (KD *>* 10,000 nM). We analyzed the modified regions as shown in Fig. A2c. The modified regions are concentrated at positions 78, 82, 83, 84, and 90. This indicates that IgGM considers these changes to be the regions that bind to Protein A. Due to the inaccuracy of existing models in predicting antigen-antibody complexes, especially for antibodies that bind in the framework region, we used AlphaFold3 to predict the structures of the single antibodies before and after the change, as shown in Fig. A2b. It can be seen from the figure that the two modified antibodies have a very slight impact on the conformation of the CDR region, almost no change. We further verified this finding through experiments. As shown in Fig. 3e, we measured the change in the binding affinity of the antibodies before and after the change to the original antigen TNF alpha. Experimental results indicate that immune affinity not only remained stable but even showed a slight improvement. This finding further supports the capability of IgGM to effectively design antibody framework regions.

To further validate the capabilities of IgGM in framework region design, we humanized the murine antibodies to mitigate the immune response induced by the framework region of murine antibodies within the human body. We have gathered five antibodies that concurrently feature murine-derived structures and humanized structures. These antibodies are designated as a test set for assessing the relevant verification indicators of humanization. To endow IgGM with the capacity for humanization, initially, we utilized BioPhi [34] to conduct the first stage of antibody humanization, thereby obtaining an initial sequence. Given that BioPhi solely took into account the sequence, its structural adaptability was not particularly high. IgGM comprehensively assimilates the structural information, ensuring that the CDR structure of the antibody remains unaltered before and after humanization. Simultaneously, it harnesses the synergistic information between the sequence and the structure to optimize the sequence. As shown below Fig. 3f, the sequence humanized by BioPhi was used to predict the structure with AlphaFold3, and the average RMSD with the existing humanized antibodies was calculated to be 0.983 Å. To ensure the retention of the humanized characteristics of the original sequence without any loss, we selected one FR region each time. Subsequently, we optimized it with IgGM. The optimized part was the portion that differed from the original murine sequence. Additionally, we implemented a frequency-based screening approach where candidate antibodies were ranked by their occurrence rate across multiple independent sampling trajectories. For the screened antibodies, we performed AlphaFold3 structure prediction and evaluated the humanization score using Abnativ. Judging from the evaluation results in Fig. 3f, in terms of the VH-Humanness Score (which quantifies sequence similarity to human germline antibodies on a 0-1 scale), the average score of IgGM designed antibodies was 0.909, significantly higher than the 0.676 score of the original murine antibodies. In each test sequence (1vfb, 4edx, 5f3b, 8euq, 4m7l), the humanization scores of IgGM designed antibodies consistently exceeded those of the corresponding murine templates. In terms of structural preservation, the average CDR3 RMSD between IgGM designed humanized antibodies and their murine templates was 0.750 Å, substantially lower than BioPhi’s 0.983 Å. Specifically, for sequences 1vfb, 4edx, and 5f3b, IgGM achieved CDR3 RMSD values of 0.691 Å, 0.736 Å, and 0.641 Å respectively. Crucially, this improved structural preservation directly correlates with functional maintenance, as evidenced by the binding affinity measurements in Fig. 3f. The lower CDR3 RMSD values indicate better preservation of the antigen-binding conformation, explaining why IgGM designed antibodies maintained binding affinity closer to the original murine antibodies compared to BioPhi-designed variants.

To further verify the effectiveness of IgGM, we conducted wet-lab verification. We used the frequency to conduct experiments on the mouse derived antibodies(PDB ID: 7K9H) [35] against the RBD of COVID-19. According to the method mentioned above, we conducted a preliminary screening based on the humanization score, and then selected 20 humanized antibodies with the highest generation frequencies across multiple sampling runs for experimental validation. After wet-lab verification, 5 humanized antibodies were obtained, as shown in Figure Fig. 3g. The affinity could be maintained at approximately the same level as the original antibody, and the affinity dissociation constant of the Mouse antibody was measured at 0.12 nM. In contrast, the affinity dissociation constants of Human-1 to Human-5 were determined to be 0.171 nM, 0.195 nM, 0.395 nM, 0.486 nM, and 0.139 nM respectively. Moreover, the humanization fractions of these human-derived antibodies all demonstrated an increase, with levels rising from the baseline mouse value of 0.657 to between 0.723 and 0.822 across the five human samples (Human-1 and Human-2 at 0.822, Human-3 and Human-4 at 0.810, and Human-5 at 0.723). This observation suggests that the humanized antibodies screened by IgGM exhibit an affinity comparable to that of the original mouse derived antibodies. The affinity values measured by Biolayer Interferometry (BLI) were plotted in Fig. 3h and Fig. A2d. The structural integrity of the designed antibodies was assessed using AlphaFold 3 predictions, with results shown in Fig. 3i. Quantitative analysis revealed an average backbone RMSD of 1.10.06 Å between the designed and original antibody structures, well below the 2.0 Å threshold typically associated with preserved functional conformation. Confirming that IgGM successfully modifies framework regions without compromising the critical antigen-binding functionality mediated by the CDR regions.

### 2.3 IgGM is an effective affinity maturation tool

IgGM can achieve the maturation of antibody affinity by only changing some amino acids. Antibody affinity maturation is a process in the immune system that gradually improves the binding ability of antibodies to antigens through mechanisms such as somatic hypermutation and clonal selection. In computer-aided antibody design, antibody affinity maturation aims to modify a known antibody to enhance its affinity. Existing methods, such as GearBind [20], regard antibody affinity maturation as a classification task. They screen for high affinity antibodies by performing single point mutations on existing antibodies and scoring them. However, such methods [20, 36] require additional input, such as the structure of antigen-antibody complexes, or additional training of a scorer for screening, are highly dependent on the effectiveness of the scoring, and involve a large search space. Selecting all possibilities is extremely time-consuming. We propose to consider affinity maturation as a generation task for a portion of the CDR region. During the generation process, the non-mutated positions remain unchanged, while the mutated positions can be customized flexibly. By combining the method of single point mutations in somatic cells, we can gradually improve the affinity. Specifically, as shown in Fig. 4a, we use IgGM as a mutator to determine the types of amino acids for single point mutations. Once the original sequence is obtained, we systematically redesign every amino-acid position within each CDR region, or alternatively pinpoint key residues manually for targeted redesign. For each position, IgGM generates 100 candidate variants through iterative sampling with different random seeds (N=50 independent sampling trajectories). Candidate antibodies are then ranked by their generation frequency, the number of independent sampling runs that produced identical sequences. This frequency-based ranking leverages the principle that sequences repeatedly generated across independent sampling trajectories represent high-probability solutions within the sequence-structure distribution learned by IgGM. After filtering out sequences identical to the original antibody, the top 10 variants with highest generation frequencies are selected for wet-lab verification. After obtaining antibodies with improved affinity, if the target affinity threshold has not been achieved, the highest-performing variant becomes the new input for the next optimization cycle. This iterative process continues until either the desired affinity is achieved or diminishing returns are observed (defined as less than 1.5-fold improvement in consecutive rounds). The iterative nature of this approach mimics natural affinity maturation while leveraging IgGM’s ability to explore sequence space more efficiently than random mutagenesis.

**Fig. 4:**
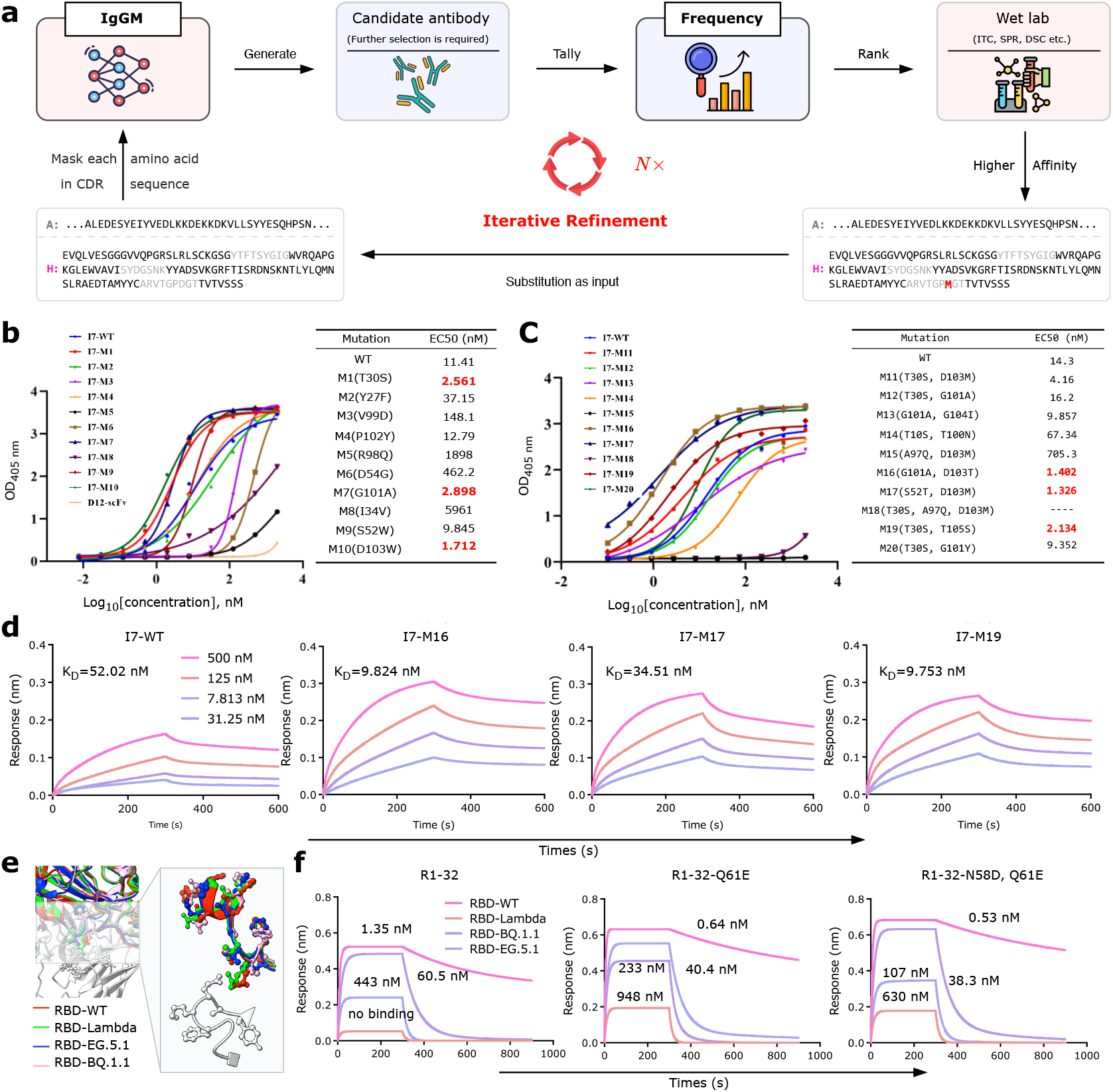
IgGM is an effective affinity maturation tool. **a**, The process of IgGM for affinity maturation. Given an antibody that needs to be matured, the amino acids of the CDRs are redesigned in sequence. Design 100 candidate antibodies for each position using IgGM. All the obtained candidate antibodies, a total of *L*_*CDR*_ *×* 100, are sorted according to the number of repetitions, that is, the frequency. The antibodies ranked at the top are selected for wet-lab verification. When an antibody with higher affinity is obtained, this antibody is used as the input for further iterative optimization. Repeat the above process until an antibody with the target affinity is obtained. **b**, I7 is an antibody that binds to human interleukin-33. We carried out two rounds of affinity optimization of I7 against IL-33. Regarding the results of the first round of affinity maturation for I7, on the left side of the figure is the measured EC50, and on the right side are the amino acids changed by mutation and their corresponding values. **c**, Regarding the results of the second round of affinity maturation for I7, on the left side of the figure is the measured EC50, and on the right side are the amino acids changed by mutation and their corresponding values. **d**, The results of the dissociation constant (*K*_*D*_) of M16, M17 and M19 selected in the second round, which were measured using BLI, showed an increase in affinity. **e**, The escape spectrum of R1-32 indicates that its affinity is highly dependent on the original conformation of RBD. Comparing the RBD structures of 7YDI (WT), 8VYG (Lambda), 8XLN (EG.5.1), and 8XYG (BQ.1.1), it can be seen that the mutation sites are concentrated on the epitope contact surface of R1-32: key residues such as L452Q/F490S (Lambda), F486P (EG.5.1), and R346T/K444T/N460K (BQ.1.1) undergo side-chain flipping or main-chain rearrangement, leading to the loss of hydrogen bonds/salt bridges and the introduction of steric hindrance. These conformational drifts prevent the CDR H3 of R1-32 from maintaining its original induced fit conformation, significantly increasing the binding free energy, ultimately resulting in a sharp decrease in affinity and complete escape. **f**, The affinity curve graphs of the two antibodies with IgGM mutation and those of the wild type. These two antibodies can function on multiple variants compared to the original antibody. At the same time, the affinity for RBD-WT is also better than that of the original antibody.

We verified the effectiveness of the IgGM based affinity maturation approach through two complementary case studies that demonstrate its versatility across different therapeutic contexts. First, we applied our method to improve the affinity of antibody I7 against human interleukin-33, representing a conventional affinity maturation scenario for a single target. Building on this validation, we then addressed the more complex challenge of developing broadly neutralizing antibodies against rapidly mutating viral targets, specifically SARS-CoV-2 variants. This second case study tests IgGM’s capability to overcome antigenic immune escape, a critical limitation of many existing therapeutic antibodies. To identify optimal mutation positions, we first analyzed the predicted antigen-antibody interface using computational alanine scanning to identify residues with high binding energy contributions. Additionally, we incorporated evolutionary conservation analysis of the CDR regions to prioritize positions with lower natural variability, which are more likely to tolerate mutations. Based on this dual criteria approach, we selected 12 key positions across the CDR regions for systematic redesign, as shown in Fig. 4a. The ELISA results of the first round are shown in Fig. 4b. The results indicate that our mutants M1, M7, and M10 achieved a 4 to 6-fold increase in affinity. Then, we carried out a new round of iteration based on these three antibodies. Since there are three antibodies with improved affinity, we generated them separately, sorted them according to frequency, and limited the number of different variants at the same time, ensuring that there are at least two variants for each of M1, M7, and M10. After that, we verified the newly selected 10 antibodies through wet experiments. The ELISA results are shown in Fig. 4c. As a result, 3 antibodies with a 5 ot 10-fold increase in affinity appeared. We further used BLI to measure the *K*_*D*_ values of different antibodies, as shown in Fig. 4d. The binding affinity improved significantly as the dissociation constant decreased from 52.02 nM for the original antibody to 9.753 nM for the best performing variant, representing a 5.3-fold enhancement in binding strength.

To further verify the effectiveness of the above process, we verified the performance of IgGM in broadly neutralizing antibody, as shown in Fig. 4e. Due to the rapid mutation of the novel coronavirus, the conformation of the RBD region may change, rendering the parts that could originally form a chemical reaction with the antibody ineffective. In order to obtain an antibody that can immunize against these variants, we used the above process to carry out affinity maturation for 4 antigens. Due to changes in the sequence structure of different variants, the distribution of the generated antibody sequences also changes. To address the challenge of viral variants, we implemented a multi-objective optimization strategy. We first performed independent affinity maturation runs for each of the four SARS-CoV-2 RBD variants (WT, Lambda, BQ.1.1, and EG.5.1). For each variant, we generated candidate antibodies and ranked them by generation frequency as previously described. We then identified the intersection set of antibodies that appeared in the top 100 candidates for all four variants, resulting in a consensus set of cross reactive candidates. From this consensus set, we selected the 20 antibodies with highest average frequency across all four variant specific optimizations for experimental validation. This approach ensures that selected candidates maintain binding capability across multiple viral variants while benefiting from IgGM’s structure aware design capabilities. At this stage, we obtained the Q61E mutant antibody. As shown in Fig. 4f, from the results, it can be seen that this mutation enabled the binding to RBD-Lambda and RBD-BQ.1.1 from no binding, but the binding affinity was still relatively weak. However, the affinity for RBD-WT was improved, proving that this mutation did not reduce the original immunity. Then, we repeated the above process based on the Q61E variant and obtained a better broadly neutralizing antibody (N58D, Q61E). As shown in Fig. 4f, especially in terms of the affinity performance with RBD-BQ.1.1, it was increased by nearly 10 times. Overall, two rounds of iteration enabled the antibody to increase its affinity for various variants while maintaining or even enhancing its affinity for RBD-WT. This demonstrates that IgGM can address antigenic immune escape and achieve broad spectrum activity, further verifying the effectiveness of the above methods.

### 2.4 IgGM can achieve *de novo* design of antibodies

IgGM enables *de novo* antibody design by generating a complete variable domain sequence directly from structural information. Given the 3D structure of an antigen, a defined epitope, and selected FRs, IgGM produces a full-length VH/VL sequence—including all six CDRs, or three CDRs in the case of nanobodies—without relying on existing antibody templates or experimental results. Starting from the designated epitope, IgGM constructs both the sequence and structure of the antibody’s variable domain, encompassing both FRs and CDRs. This approach enables the generation of antibodies with high affinity and precise epitope specificity. To ensure better properties, the framework regions input to IgGM can be selected based on any favorable biophysical properties. As illustrated in Fig. 5a, IgGM supports iterative generation of antibody sequences and structures. For further details on framework selection, refer to the non-colored section of Fig. A3f.

**Fig. 5:**
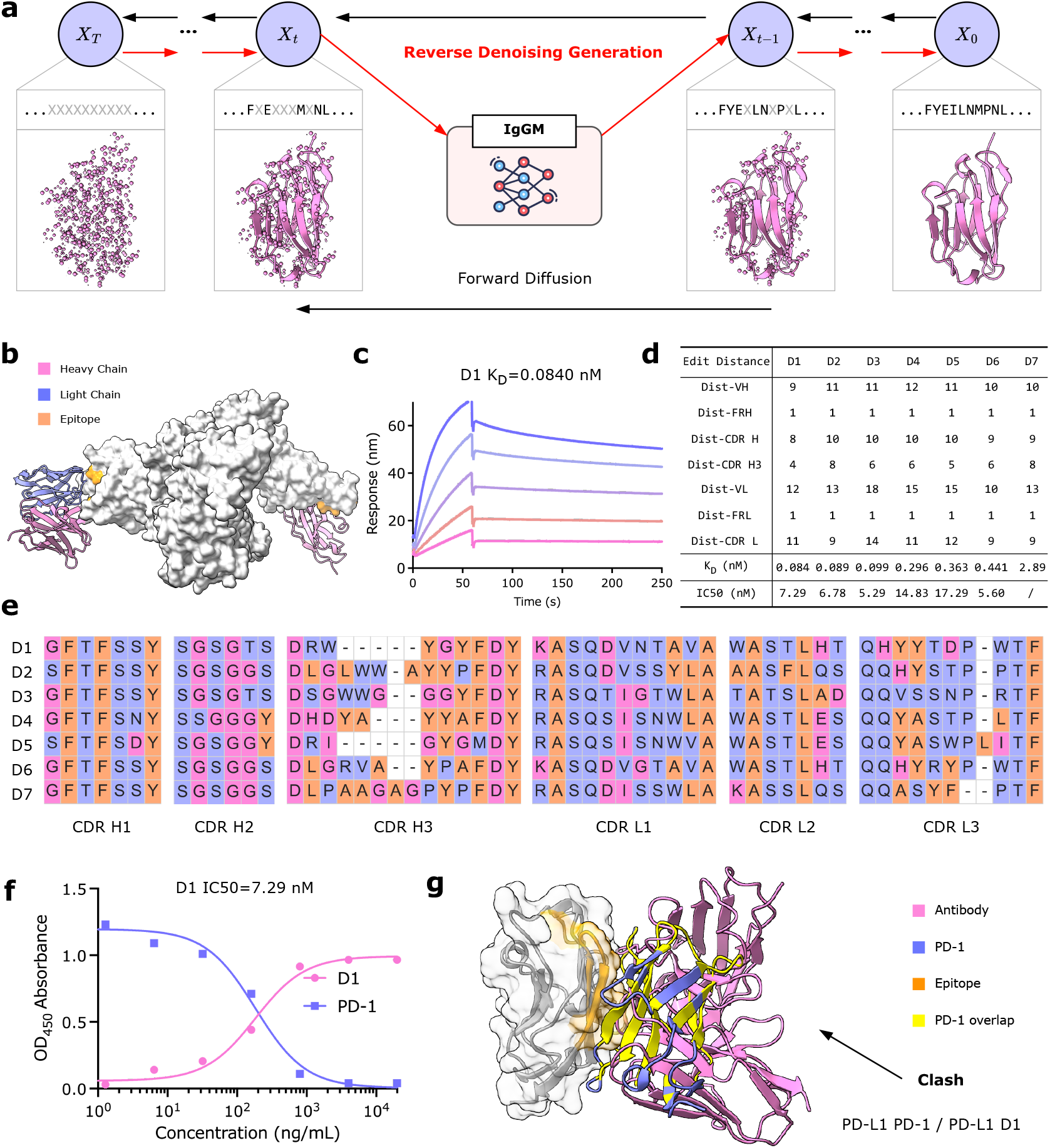
IgGM can achieve *de novo* design of antibodies. **a**, IgGM can generate the sequences and structures of antibodies through a multi-step iterative process. **b**, IgGM can generate corresponding antibodies by targeting specific epitopes. The figure shows the antibodies and nanobodies designed by IgGM against different epitopes. **c**, The best bindable antibody designed by using IgGM against the PDL1 target can achieve a maximum *K*_*D*_ of 0.084 nM. **d**, Results showcasing the affinity and novelty of the best antibodies designed by IgGM for PD-L1. Affinity is evaluated using binding KD through SPR. Novelty is measured using the edit distance to the closest antibody in various regions of the antibody sequence. **e**, CDR multiple sequence alignment of seven antibodies designed by IgGM, highlighting significant diversity. **f**. Results of ELISA competitive binding assay. D1 blocks the PD-1/PD-L1 interaction in a dose-dependent manner. **g**. Predicted structural models indicate that our *de novo* antibodies (D1, consisting of heavy and light chains) effectively block the interaction between PD-1 and PD-L1 (PDB ID: 3BIK, which includes PD-1 and PD-L1). The overlapping parts of PD-1 with the antibody are highlighted in yellow, with a calculation threshold of 1 angstrom, and there are numerous clashing parts.

To benchmark IgGM *in silico*, we evaluated its ability to recover CDR sequences and structures across 60 antibody–antigen complexes from the SAb23H2 test set, derived from the SAbDab database [29]. For dataset details, see Section 4.1. We compared four generative models: MEAN [37], DiffAb [38], dyMEAN [30], and IgGM, using amino acid recovery rate (AAR) for sequence accuracy and backbone RMSD (N, C_*α*_, C, O) for structural fidelity. Since MEAN and DiffAb do not support sequence design outside the CDR regions, we used IgFold [13] and AlphaFold3 [11] to initialize antibody backbones and generate CDR sequences. These structure prediction tools leverage advanced algorithms to model protein structures from sequence inputs, enabling accurate structure modeling. MEAN is limited to CDR H3, so its metrics are reported only for that loop, while the other models generate all six CDRs simultaneously. Across all CDRs, IgGM consistently achieved the highest AAR and lowest RMSD (Fig. A3a, b). Notably, for the highly flexible CDR H3 loop, IgGM reached 36% AAR—an absolute improvement of 22.4% over the previous best (dyMEAN, 13.6%). Additionally, we found that using AlphaFold3-predicted structures as initialization inputs led to more realistic antibody conformations, further enhancing design quality. In addition, unlike existing methods [37, 38], which require strict structural constraints on known antigen-antibody complexes, IgGM is capable of generating antibodies for antigens without any known binding partners. By specifying the epitope location, IgGM can produce antibodies that target precise spatial positions, as illustrated in Fig. 5b. Furthermore, IgGM supports the customization of CDR region lengths, enabling flexible design tailored to specific structural requirements. This capability is applicable to real-world antibody engineering scenarios, as demonstrated in Fig. A3c.

While *in silico* evaluations confirm IgGM’s ability to generate structurally accurate antibody designs, the ultimate validation lies in experimental testing of binding affinity and specificity. To assess IgGM’s practical utility in clinically relevant antibody development, we conducted a comprehensive wet-lab study targeting Programmed death-ligand 1 (PD-L1), a key immune checkpoint protein and established target in cancer immunotherapy. Leveraging IgGM’s ability to incorporate customized framework regions, we selected FRs from specific humanized V and J genes—namely IGHV323/IGKV15 and IGHJ4/IGKJ1—as input templates. For the antibody CDR regions, the lengths are variable, particularly for CDR3. According to statistical analysis by Oscar et al. [39], CDR3 exhibits the greatest length diversity. We visualized the CDR H3 length distribution in Fig. A4a. In contrast, CDR1 and CDR2 are determined by the V gene and have fixed lengths (H1: 7, H2: 6, L1: 11, L2: 7). To determine appropriate CDR3 lengths, we analyzed antibodies sharing the same V gene in the OAS database and selected the most frequent combinations—7 to 18 residues for CDR H3 and 9 to 11 for CDR L3. To further constrain the design space, we integrated the statistical distribution from [39] and narrowed the CDR H3 length range to 8–15 residues, balancing diversity with structural feasibility. We designated the binding sites of Programmed cell death protein 1 (PD-1) and Programmed death-ligand 1 (PD-L1) as the epitope. According to the results presented in [22], the experimental success rate of *de novo* antibodies is very low. This is because the structure prediction method performs poorly in predicting the CDRs of antibodies. To avoid the cumulative errors caused by relying on other methods, we designed a comprehensive experimental validation pipeline. First, based on our experience in affinity maturation, we designed for different lengths. For the different lengths of CDR H3 and CDR L3, we conducted random length combinations, obtaining 24 length combinations here. Then, for these 24 length combinations, we used IgGM to generate 10,000 candidate antibodies. We computed the edit distance between each candidate antibody sequence and all known positive antibodies in public databases (SAbDab), removing candidates with an edit distance of less than 5 from any known sequence. CDR sequences differ by more than five amino acids from any known antibody are enriched for novel paratopes and are more likely to exhibit distinct binding modes. Next, we performed further filtering based on frequency and the confidence of AlphaFold 3-predicted structures [11], ultimately selecting the top 60 sequences ranked by occurrence for wet-lab experimentation. After wet-lab experiment results, this final set included seven antibodies with nanomolar or even picomolar affinity, calculated according to the KD value, representing a 7/60 success rate among the high affinity candidates. The final characterization identified seven high affinity antibodies with KD values ranging from 0.084 nM to 2.89 nM, all demonstrating specific binding to PD-L1.

The 7 lead antibodies are shown in Fig. 5c and Fig. A3e. Among them, D1 showed an affinity of 0.084 nM, Fig. 5e illustrates the CDRs of the 7 PD-L1 antibodies. We observed that while CDR-H1 and CDR-H2 displayed moderate similarity (dictated by the V gene), CDR H3 and CDR L3 demonstrated two distinct diversities. This combinatorial diversity in CDR regions dramatically expands the druggable sequence space [40], enabling systematic exploration of non-immunogenic paratopes that would otherwise remain inaccessible through conventional immunization or library screening methods. To further verify the effectiveness of the generated antibody, we performed competition assays against PD-1 on seven antibodies using ELISA. As shown in Fig. 5f, with the increase in D1 concentration, the D1-PD-L1 binding signal (red curve) gradually increased, while the PD-1-PD-L1 binding signal (blue curve) decreased accordingly. The results of D2-D6 can be found in Fig. A3f, with a blocking rate of 6/7 achieved, indicating that all these PD-L1 antibodies bind to the PD-1 epitope-specific. Consistently, D1 displaced PD-1 from PD-L1 in a concentration dependent fashion. D1 effectively prevented PD-1 ligation. Specifically, the ELISA measurements for D1 highlighted its competitive binding capabilities: D1 showed substantial binding to PD-L1 alone, yet its RU values markedly declined when combined with PD-1, demonstrating its competition with PD-1 for PD-L1 binding. The results of aligning the IgGM generation structure with the structures of PD-1 and PD-L1 are presented in Fig. 5g. There is a significant structural clash between D1 and PD-1. The predicted structure indicates that D1 can effectively bind near the epitope of PD-L1, thereby blocking the interaction between PD-1 and PD-L1. Structural analysis revealed the precise mechanism of competitive inhibition. A superposition of the IgGM predicted D1–PD-L1 complex with the crystal structure of PD-1–PD-L1 (PDB 3BIK) shows a 7.3 Å steric clash between the D1 heavy-chain CDR H3 and PD-1 (Fig. 5g). This structural interference occurs precisely at the PD-1/PD-L1 interface, with the D1 CDR H3 occupying the same spatial region as PD-1’s binding loop. The magnitude of this clash (7.3 Å) substantially exceeds typical van der Waals overlap thresholds (typically 1-2 Å), confirming that D1 binding would physically prevent PD-1 engagement with PD-L1. This structural insight directly explains the competitive binding observed in our ELISA assays (Figure Fig. 5f), where D1 concentration-dependently displaced PD-1 from PD-L1.

## 3 Discussion

Our study demonstrates the feasibility of unifying multiple antibody design tasks within a single model, spanning *de novo* antibody generation, affinity maturation, framework engineering, structure prediction, and inverse design. Experimental validation against diverse antigens—including PD-L1, Protein A, TNF-*α*, IL-33, and the SARS-CoV-2 RBD with its variants. In addition, we participated in AIntibody competition [41] with IgGM, achieving the optimization of pM-level SARS-CoV-2 RBD antibodies and earning a top-three ranking. This confirms IgGM’s ability to generate high-affinity antibodies and nanobodies, underscoring its broad applicability.

Unlike traditional computational approaches that depend on prior structural templates [37, 38] or generate large libraries followed by downstream filtering [22, 42], IgGM is trained extensively on specific antigen–antibody complexes, enabling it to directly learn the principles of antigen–antibody recognition. This endows the model with the capacity to maximise the generation probability of functional binders, inspiring a frequency-based selection strategy that has been validated across tasks, and highlighting the centrality of modelling antigen–antibody interactions in effective antibody design.

Despite these advances, limitations remain. IgGM relies on fixed antigen and epitope inputs, precluding it from capturing binding-induced conformational dynamics—a gap that could be addressed by incorporating structural kinetics. Its current evaluation is limited to single antigens, with performance on complex antigens such as pMHC yet to be established. Furthermore, IgGM models only backbone atoms, omitting explicit side-chain representation that is critical for forming functional sites, driving hydrophobic packing, and conferring binding specificity [43]; incorporating side-chain modelling, as in full-atom generative frameworks [11], offers a promising direction.

By jointly modeling antigen–antibody interactions and sequence–structure correspondence, IgGM advances towards a unified framework for antibody engineering challenges, mirroring the broader shift in other domains from task-specific methods to general-purpose generative models [44]. Decoupling antibody design from template dependency opens new possibilities for targeting “undruggable” epitopes lacking natural binders. Moreover, the proposed frequency-based prioritisation strategy illustrates that generative models can serve not only as design engines but also as intrinsic ranking mechanisms, enabling more rapid and cost-effective candidate selection. Collectively, IgGM offers a new paradigm for AI-driven antibody design—capable of continuously optimising antibodies across development stages—and substantially expands the application scope of computational antibody engineering.

## 4 Methods

### 4.1 Dataset

We constructed our training, validation, and test sets from the SAbDab database, employing the widely used method of dividing the dataset based on time, as previously established in other works [15, 45–47]. We selected all experimentally determined antibody structures published in the database up to December 31, 2022, as our training set. The final training set consisted of 6,448 antibody-antigen complexes with both heavy and light chains and 1,907 single-chain antibody-antigen complexes. During the training process, we used CD-Hit [48] to cluster the training set, with each cluster containing antibodies with sequence similarities above 95%, resulting in a total of 2,436 clusters. To ensure the utilization of available data, we randomly sampled one sample from each cluster for training in each epoch. The validation and test sets included antigen-antibody complexes determined experimentally and published between January 1, 2023, and June 30, 2023, and between June 30, 2023, and December 30, 2023, respectively. We removed samples with sequence similarities above 95% to those in the training set to eliminate redundancy in the data, ensuring a fair evaluation. This process resulted in 101 and 60 validation and test set samples, respectively, which were completely unrelated to the training set. The validation set was used for hyperparameter tuning and model selection, while the test set was used for the final model evaluation. We named this test set SAb23H2. The specific data processing methods can be found in the Appendix D.1.

### 4.2 Denoising diffusion probabilistic models

Diffusion models gradually add noise to the data through a forward process until it becomes random noise, and then learn a reversible backward process to progressively recover the original data from the noise. Training is achieved by minimizing the KL divergence between the forward and backward processes. When the backward process is parameterized in a specific way, the model is equivalent to multi-scale denoising score matching and annealed Langevin dynamics, capable of generating high-quality samples. We utilized the framework of diffusion models to co-design the sequences and structures of antibodies.

In our study, we represented the sequence information of antibody-antigen complexes through amino acid sequences and described the structure of proteins and the orientation of each amino acid using the coordinates of the *C*_*α*_ and the relative positions between backbone atoms, with the generation of these representations facilitated by diffusion models. By extracting the three-dimensional coordinates of the *C*_*α*_ of each amino acid, we constructed a simplified model of the protein structure that retained the structural

features of the backbone of the protein [38, 49]. Additionally, in order to more accurately represent the orientation of amino acids in space, we calculated the relative positions between the backbone atoms of adjacent amino acids, thus obtaining a directional framework that described the protein structure. This modeling approach effectively preserved key information of the protein structure, providing theoretical support for subsequent structural design. Inspired by previous work [50–53], we employed a hybrid diffusion model for the generation of antibody sequences and structures.

For antibody sequences, we employ a discrete diffusion model for generation. Specifically, there are 20 classes of amino acids. For a specific antibody sequence, the amino acid type at each position can be regarded as a categorical distribution. For a random variable at a certain position, there are 20 classes, denoted as *s*_*t*_, …, *s*_*t−*1_ *∈ {*1, …, 20*}*, the forward noise-adding process can be written as:

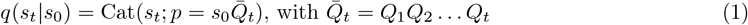

We can obtain the following *t*-step marginal and posterior at time *t −* 1:

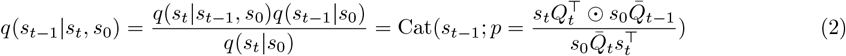

where *s*_0_ is the original data, Cat(*s*; *p*) represents a categorical distribution, and *s* is a 20-dimensional one-hot vector indicating the probability of a specific class of amino acid occurring at that position, denotes the element-wise product.

For the *C*_*α*_ coordinates of antibodies, the values in three-dimensional space are continuous, and therefore a continuous diffusion model can be used to generate the *C*_*α*_ coordinates [49]. In practical applications, Gaussian noise can be gradually added to the coordinate values until they approach a Gaussian distribution. Specifically, for the coordinate **x**_*t*_, the process of adding noise in the forward process can be written as:

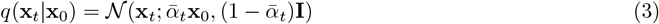

where 𝒩 (**x**_*t*_; *µ, σ*^2^) represents a normal distribution with mean *µ* and variance *σ*^2^, 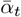 is a scalar less than 1. We can obtain the following *t*-step marginal and posterior at time *t −* 1:

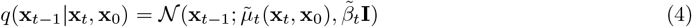

where 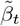 is about the cumulative product of the variance schedule up to time step *t*.

For each amino acid direction, the orientation within the local coordinate system of each amino acid can be considered as a continuous value in the SO(3) space, analogous to a uniform distribution on polar coordinates [52]. Consequently, we can construct both forward and backward diffusion processes on the three-dimensional rotation group SO(3). The forward process diffusing the direction data *o*_0_ into pure noise, following the specific formula:

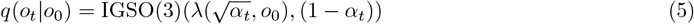

where IGSO(3) denotes the isotropic Gaussian distribution on SO(3), and *λ* represents the scalar product along the geodesic from the identity rotation matrix to *o*_0_. Conversely, the backward process is designed to transform noise back into data, guided by the following probability distribution:

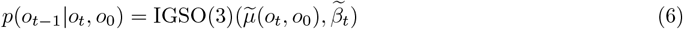

where 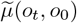 denotes the mean of the backward process, which is calculated as the product of two rotation matrices.

### 4.3 Feature extraction from protein language models

Inspired by the success of pre-trained language models in natural language processing, such as BERT [54] and BART [55], the bioinformatics and computational biology communities have recently turned their attention to pre-trained protein language models [56–58]. These innovative models conceptualize protein sequences as a unique language leveraging masked language modeling techniques for their training. Protein language models are generally built upon the Transformer architecture, which is particularly adept at handling extensive sequence data. Through the process of training on vast protein sequence datasets, these models acquire a deep understanding of the syntactic and semantic elements inherent in protein sequences. The utility of such large-scale language models in the biological domain has been demonstrated by a multitude of studies [56, 59–62].

Given that antibody-antigen complexes are composed of multiple chains, we have selected Pre-trained Protein Sequence Model(PPSM [15]) as our sequence feature extractor due to its ability to handle inter-chain relationships. PPSM is an extension of the ESM2 model [56], which has been further refined to improve its proficiency in capturing the structural and functional characteristics of multi-chain protein complexes. The enhanced capabilities of PPSM for multi-chain protein structure prediction are detailed in [15], where readers can find a more in-depth discussion of the model’s architecture, training procedure, and its application in the context of our research.

### 4.4 Denoising network architecture

To facilitate the recovery of antibody sequences and structures from perturbed data, we introduce a architecture that leverages a pre-trained protein language models(PLM). The antigen and perturbed antibody data are processed by the PLM, with the features from the final layer being meticulously extracted to serve as input to our denoising network. To preserve the integrity of the learned features and to conserve computational resources, we maintain the PLM’s parameters in a frozen state throughout this process. To enhance the model’s ability to discriminate between different chains within the antibody, we incorporate chain-specific representations into the PLM’s output features. This allows the model to develop the understanding of the distinct chains that comprise the antibody structure. Furthermore, to address the critical aspect of antigen epitopes, we augment the antigen feature set with specialized representations that highlight the interactions between the antibody and the antigen. In order to provide the model with a means to identify the precise positions of individual amino acids, we employ a difference sine encoding scheme. This scheme codes for both the temporal steps and the sequence positions, facilitating the model’s recognition of the spatial organization of amino acids within the antibody sequence.

The backbone of our model is meticulously designed to integrate 16 layers of Sgformer modules with 8 layers of Predict modules, creating a powerful framework for the prediction and refinement of antibody sequences and structures as shown in Fig. 1c. The Sgformer module serve as the initial processing stage, where they receive the feature provided by the PLM. These modules are designed to distill the essence of the antibody sequence through their multi-layered architecture, facilitating the learning of sequence features and pairwise interactions. The sequence features extracted at this stage are pivotal for the subsequent recovery of the original sequence from the perturbed input, while the pair-wise representations encode the relational information essential for understanding the complex folding of the antibody-antigen complex. ransitioning from sequence to structure, the Predict modules play a crucial role in refining the spatial configuration of antibody models. We employ an SE(3)-equivariant invariant point attention mechanism [45], which ensures that predictions remain consistent regardless of the antibody’s orientation or position in 3D space. This equivariance is essential for accurate structural reconstruction, as it enables the model to focus on intrinsic geometric relationships between amino acids, rather than being influenced by extrinsic transformations such as rotation or translation. The Predict modules take full advantage of the sequence features and pair-wise representations learned by the Sgformer modules. By integrating these features with the invariant point attention mechanism, the Predict modules are able to iteratively refine the coordinates of the amino acids, ultimately revealing the precise spatial arrangement of the antibody’s three-dimensional structure.

### 4.5 Loss function

In the context of diffusion models, the objective can be either to predict the added noise or to recover the original target [51], more detail can be found in Section D.3. For our hybrid diffusion model, we have chosen to design our functions with the goal of recovering the original target. Inspired by previous work, we have established a suite of loss functions to train our model to achieve the desired recovery. For amino acid sequences, given that there are only 20 types of amino acids, moreover, different amino acids exhibit similar backbone atoms. We treat sequence recovery as a classification problem. We employ cross-entropy loss to guide the model in learning the correct sequence.

When it comes to the structure of antibodies, as previously mentioned in Section 4.2, our objective is to recover the spatial coordinates of alpha carbons and the orientations of backbone atoms. To this end, we utilize the residue Frame Mean Squared Error (FMSE) loss, which has been demonstrated to be effective in [49]. This loss function is specifically designed to measure the discrepancy between the predicted and actual frames of protein residues, which are essential for accurately modeling the three-dimensional structure of antibodies. We have further developed an enhanced loss function termed inter-chain Frame Mean Squared Error (iFMSE) loss. This loss function is specifically tailored to impose constraints on the differences between different chains within the antibody structure. The iFMSE loss is designed to ensure that the model accurately captures the relative orientations and positions of the various chains that make up the antibody, which is crucial for maintaining the integrity of the quaternary structure. Additionally, we employ the inter-residue distance and angle metrics to reconstruct the orientations of backbone atoms. By incorporating these loss functions, our model is trained to not only accurately predict the amino acid sequences but also to reconstruct the intricate three-dimensional structures of antibodies, thereby enhancing the potential of our approach in the fields of structural biology and antibody modeling. More detail about loss function can be found at Section D.3.

### 4.6 Evaluation criteria for antibody design

For the sequence portion, we employ metrics that have been widely used in previous work [21, 30, 38, 63, 64]: AAR, the sequence recovery rate represents the proportion of similarity between the designed antibody sequence and the actual antibody sequence. A higher value indicates that the model has a greater ability to generate a specific antibody.

For the evaluation of antibody structure design, we employ several established protein structure assessment metrics, including the RMSD, TM-Score [27], GDT-TS [65], DockQ [28], and SR.

- TM-Score (Template Modeling Score): This metric measures the structural similarity between the predicted antibody structure and a reference structure. A TM-Score of 1 indicates an exact match between the two structures, while a score closer to 0 indicates a poor match, a high TM-Score suggests that the predicted structure closely resembles the native structure.
- GDT-TS (Global Distance Test-Total Score): This metric provides an overall assessment of the model’s accuracy by comparing the predicted structure to the native structure based on the global distance test. It takes into account both the accuracy of the model’s predictions and the conformational similarity to the native structure. A higher GDT-TS score indicates a better match between the predicted structure and the native structure, suggesting a higher quality design.
- DockQ: This metric is specifically designed for evaluating the quality of protein-protein docking predictions. It assesses the interface complementarity and the conformational accuracy of the predicted complex structure. A high DockQ value indicates that the predicted interface is likely to be functional and stable, suggesting a well-designed multi-chain interface.

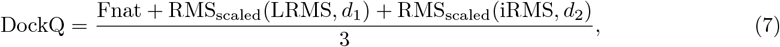

where RMS_scaled_ represents the scaled RMS deviation corresponding to either LRMS or iRMS, *d*_*i*_ is a scaling factor, *d*_1_ is used for LRMS, and *d*_2_ is used for iRMS. Fnat is defined as the fraction of native contacts retained in the predicted complex interface.
- SR (Success Rate): Indicates that the quality of the designed multi-chain interface positioning is within an acceptable range when DockQ is greater than 0.23. High SR indicates that more structure of antibodies is good.

#### Algorithm 1 IgGM Sampling

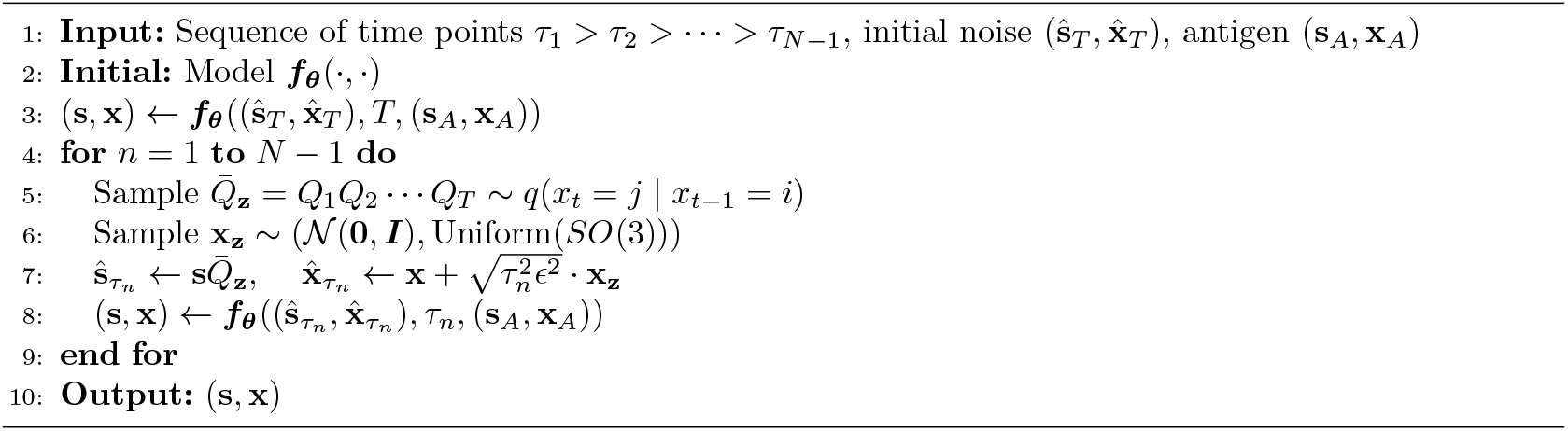

### 4.7 Training procedure and Generating procedure

The training process of IgGM is divided into two stages, following the concept of curriculum learning. Initially, we train the model for structural design. Given an antigen-antibody complex, we perturb the antibody structure to introduce noise, and then have the model reconstruct the perturbed antibody structure. During this process, we employ self-conditioning [66] to enhance the stability of the training. This self-conditioning technique involves feeding the model with additional information derived from the original unperturbed structure, which helps the model to better learn the underlying patterns and regularities in the data. Once the first-stage model has converged, we use the parameters from the first-stage training to proceed with the second stage. In the second stage, we perturb the sequence and structure of the antibody’s CDR regions and have the model reconstruct the perturbed antibody. This perturbation is aimed at introducing greater complexity and variability into the training data, thereby challenging the model to generalize better to unseen data. Throughout both stages, we use self-conditioning to enhance the training stability. This approach ensures that the model learns robustly and can generalize well to new and challenging data. The two-stage training process allows the model to first learn the basic patterns and then progressively build upon that knowledge to handle more complex scenarios. The specific training details can be found in the Section D.4. During the generation process, we adopted a consistency model [67] to accelerate the sampling speed for antibody sequences and structures, as traditional diffusion models often exhibit sluggish performance. The sampling procedure is detailed in Algorithm 1. The consistency model allows for the instantaneous generation of the final result in a single step. Alternatively, the model can be optimized through multiple iterations to enhance the stability of the generated results. The effects of different step numbers on the outcomes are discussed in the appendix, which provides a comprehensive analysis of the consistency model’s performance under various sampling strategies.

## Data Availability

All data is freely available from public sources. Experimental structures are drawn from the same copy of the SAbDab [29] at https://opig.stats.ox.ac.uk/webapps/sabdab-sabpred/sabdab.

## Code Availability

The source code, weights and inference scripts for the IgGM models are available at https://github.com/TencentAI4S/IgGM.

## Acknowledgements

This work was supported in part by Science and Technology Innovation (STI) 2030—Major Projects under Grant 2022ZD0208700, and National Natural Science Foundation of China under Grant 62376264.

## Author contribution

R.W., F.W., P.Z., X.G. and J.Y. conceived the study. R.W. developed the code and trained IgGM with the assistance of F.W. . R.W. generated experimentally characterized designs. J.S., Y.K. and T.Y. conduct wetlab validation experiments of Protein A and IL-33. J.M. assisted in the experiments related to humanization. Y. S., B.H. and Q. Y. conducted a wet experimental verification of broad-spectrum antibodies. R.W. and F.W. wrote the manuscript. R.W. drew the pictures in the manuscript. All authors read and contributed to the manuscript.

## A Extended Data Figures

**Fig. A1:**
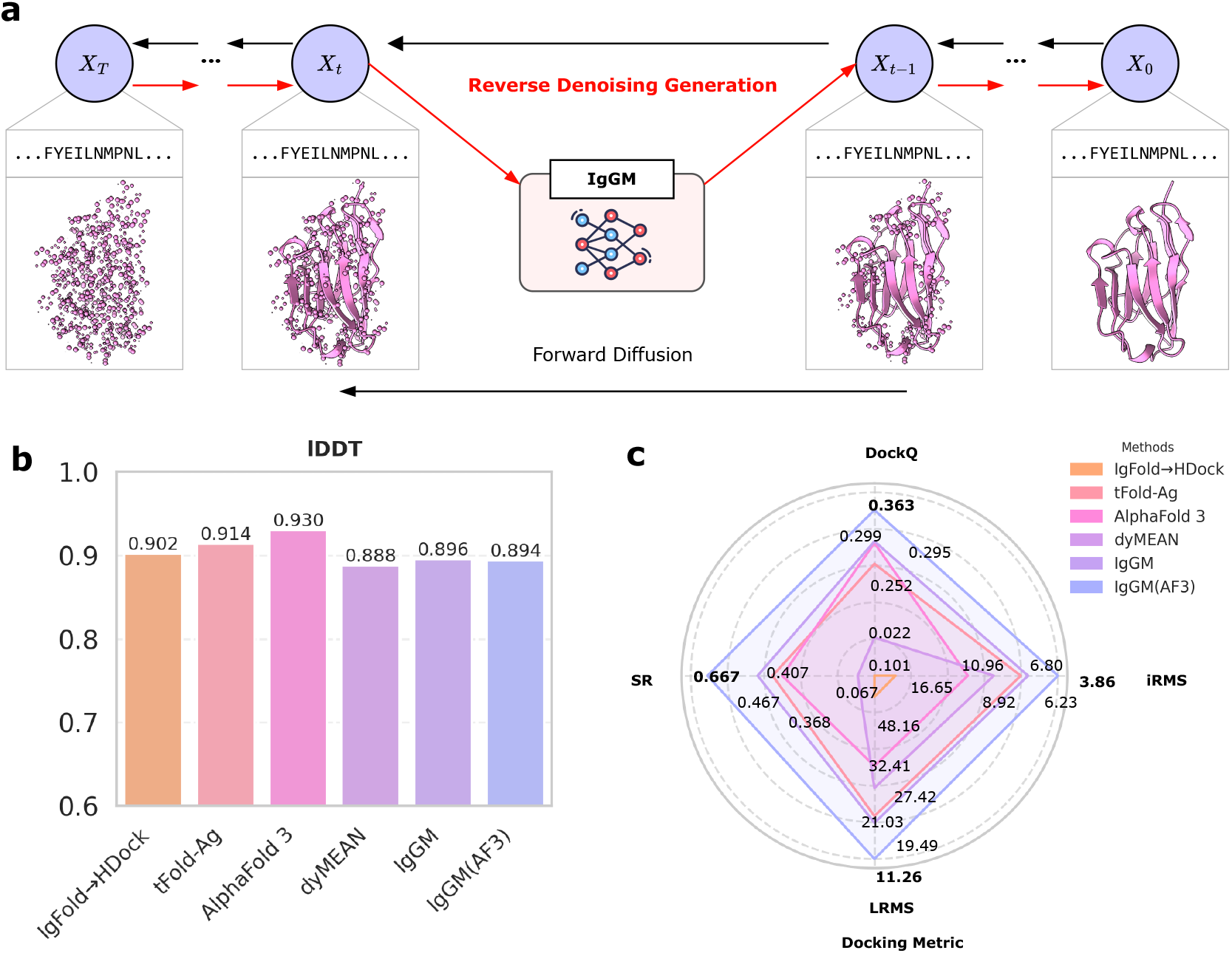
**a**, Flowchart of antibody structure prediction. The entire inverse design only involves changes in the structure. **b**, Performance of the structure prediction method on antibody structures lDDT. IgGM can almost match the most advanced protein structure prediction models. **c**, Performance of the structure prediction method on complex structures. IgGM can outperform other methods and achieve the highest prediction performance.

**Fig. A2:**
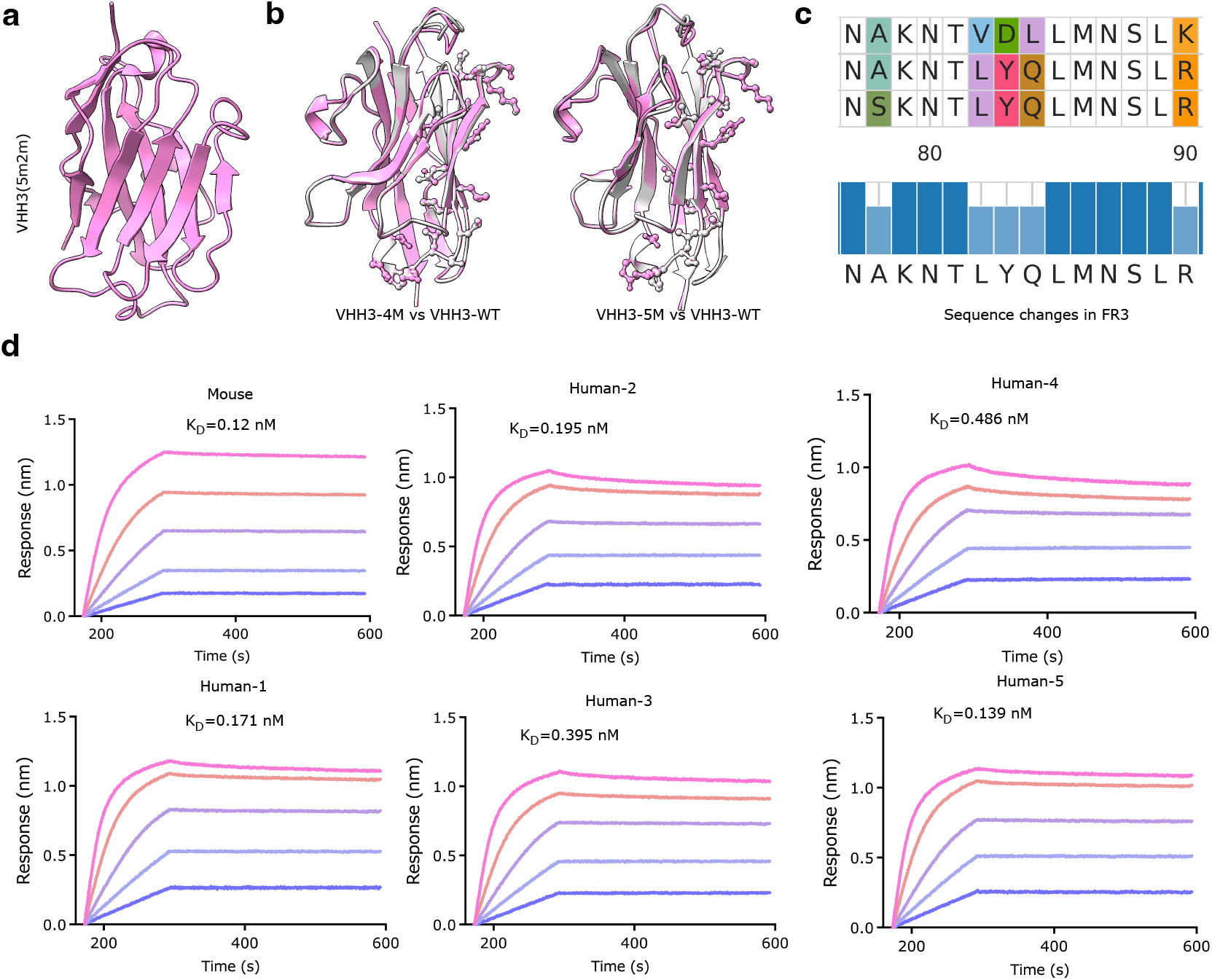
**a**, The structure of VHH3. VHH3 can bind to human TNF alpha, but the original framework cannot bind to Protein A. **b**. Use AlphaFold 3 to predict the structures of the antibody before and after the changes. The changes of 4M and 5M both bring about changes in the side chains of the framework region. **c**. Sequence comparison of VHH3-4M, VHH3-5M and VHH3-WT on FR3. 4M changed 4 amino acids, and 5M changed 5 amino acids. **d**. All affinity curve graphs of human-derived antibodies and mouse derived antibodies.

**Fig. A3:**
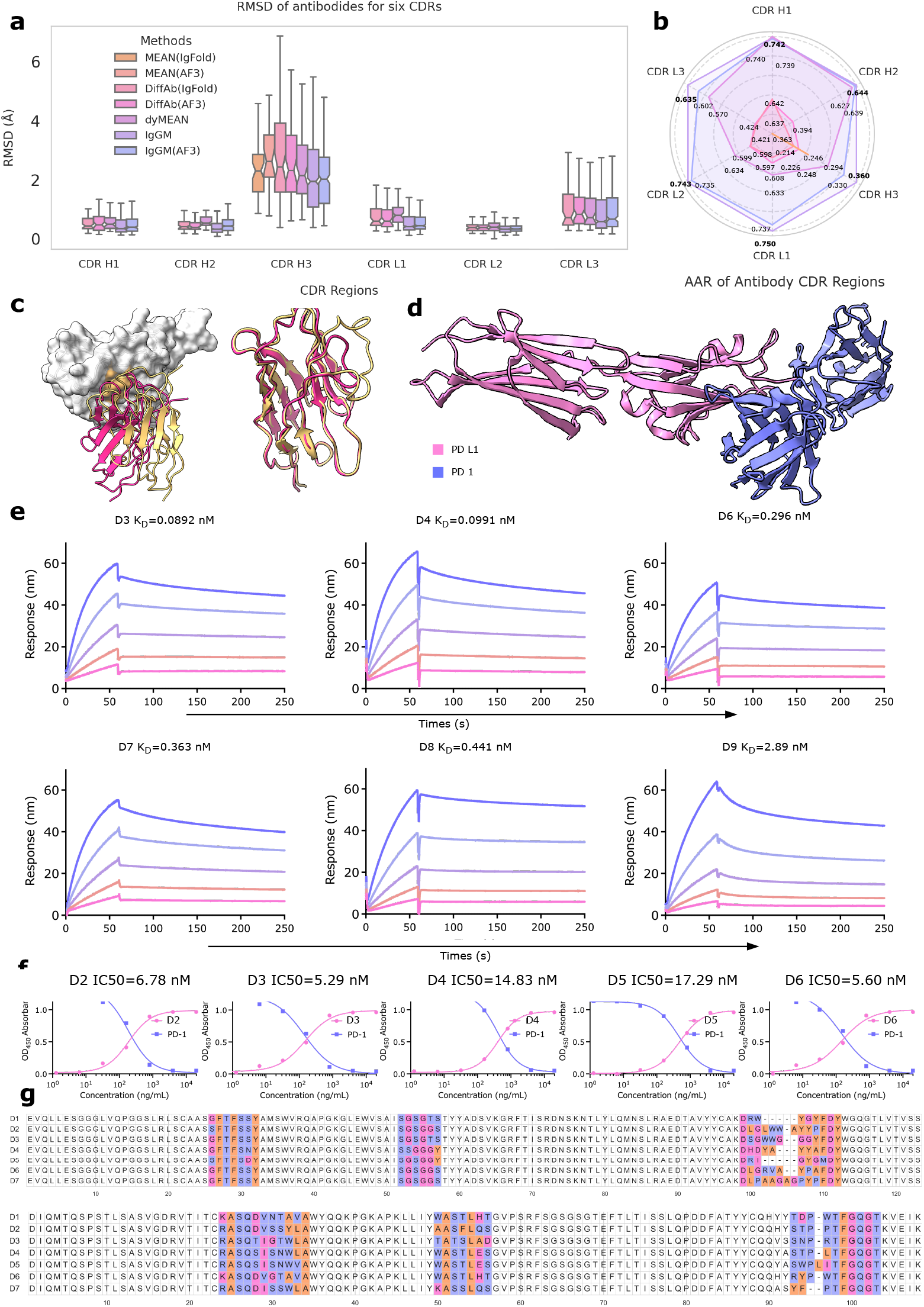
**a**, The distribution diagram of RMSD, by measuring the root mean square deviation of the backbone heavy atoms (N, C_*α*_, C, O) between the predicted and the experimentally determined structures **b**, The sequence recovery rates in different regions. Different colors represent different methods. The larger each axis is, the higher the sequence recovery rate indicates. **c**, IgGM can design corresponding antibodies by specifying the length of the CDR region. The left side of the figure shows a nanobody with CDR3 lengths of 10 and 20; both bind to the epitope, and the conformation of CDR3 will change. **d**, The binding structure of PD-L1 and PD-1. **e**, Binding kinetics of four antibody variants (D2-D7) to PD-L1, measured by surface plasmon resonance (SPR). Affinity constants range from 0.084 nM to 2.89 nM. **f**, Sequence alignment of variable regions for D1-D7, highlighting divergent residues. The CDR H3 region shows obvious differences.

**Fig. A4:**
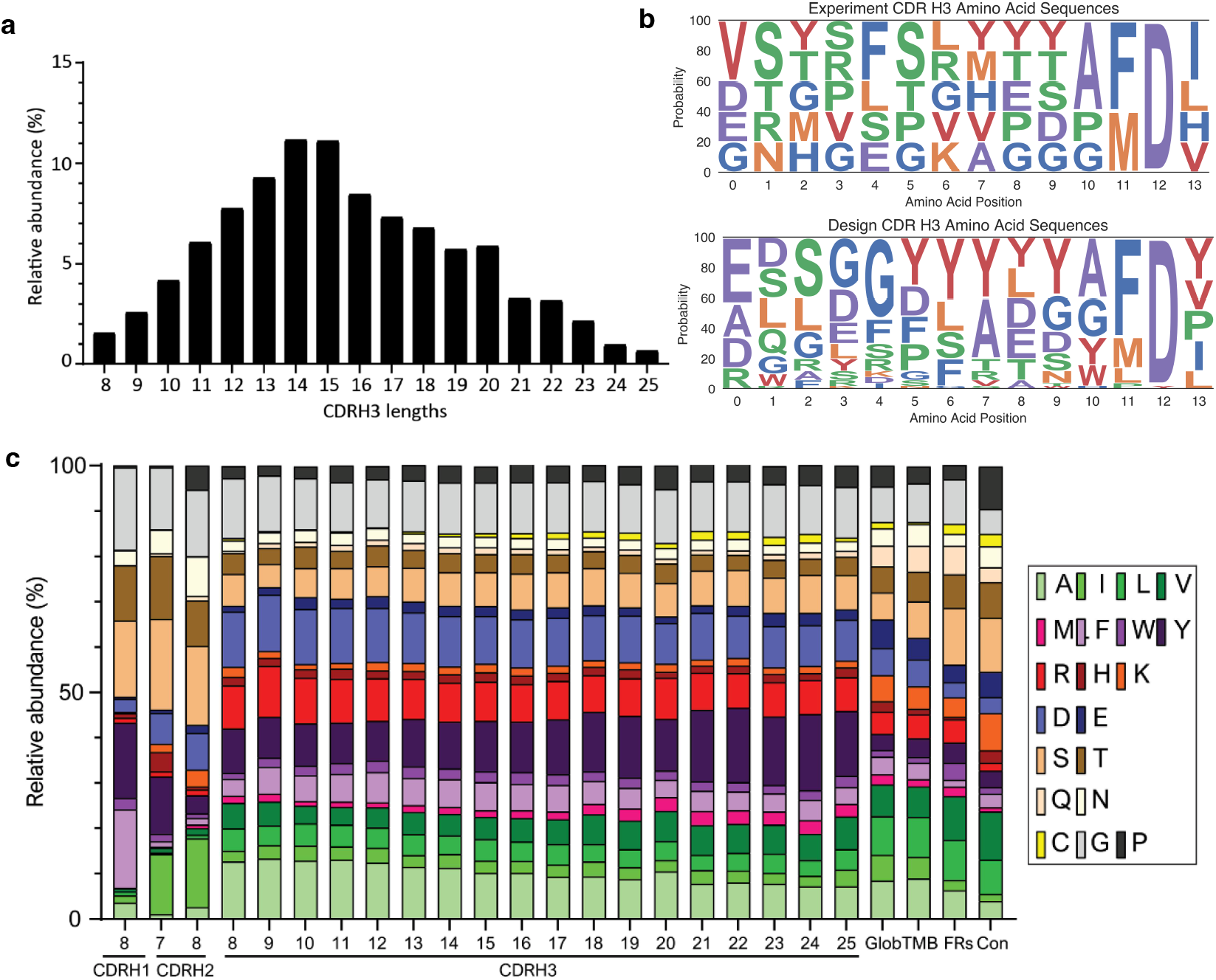
Some amino acid sequence statistics. The statistics for (a) and (b) are derived from [39]. **(a)** The length distribution of CDR H3 in antibodies. **(b)** Logo plot of the CDR H3 sequence distribution designed by IgGM, with sequences of length 15. **(c)** The proportion of different types of amino acid sequences contained within CDRs of varying lengths.

**Fig. A5:**
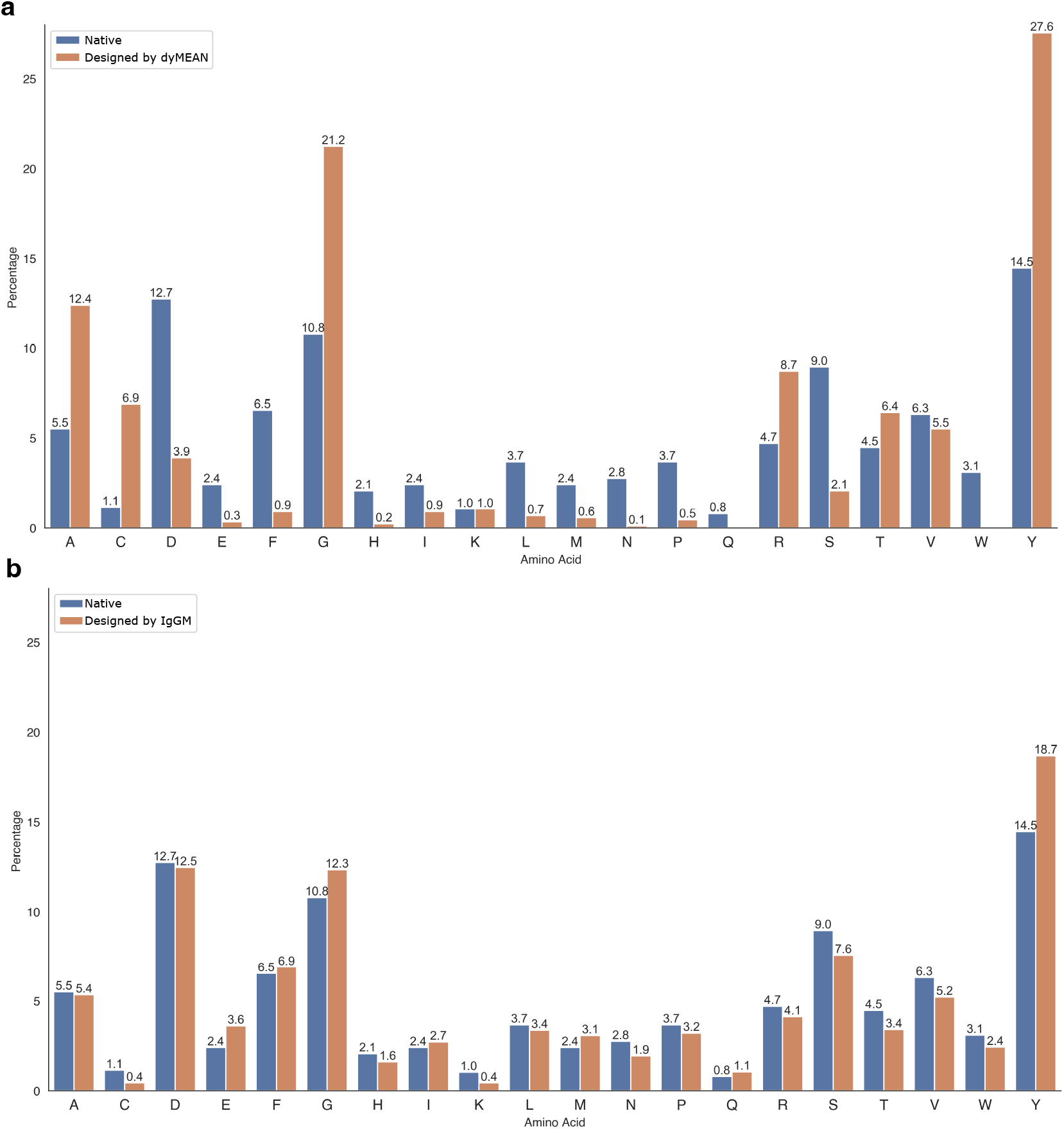
Statistics of the probabilities of generating amino acids in different CDR regions. **(a)** The results generated by dyMEAN mostly concentrate on G and Y, deviating from the native distribution. **(b)** The results generated by IgGM are more in line with the native distribution.

**Fig. A6:**
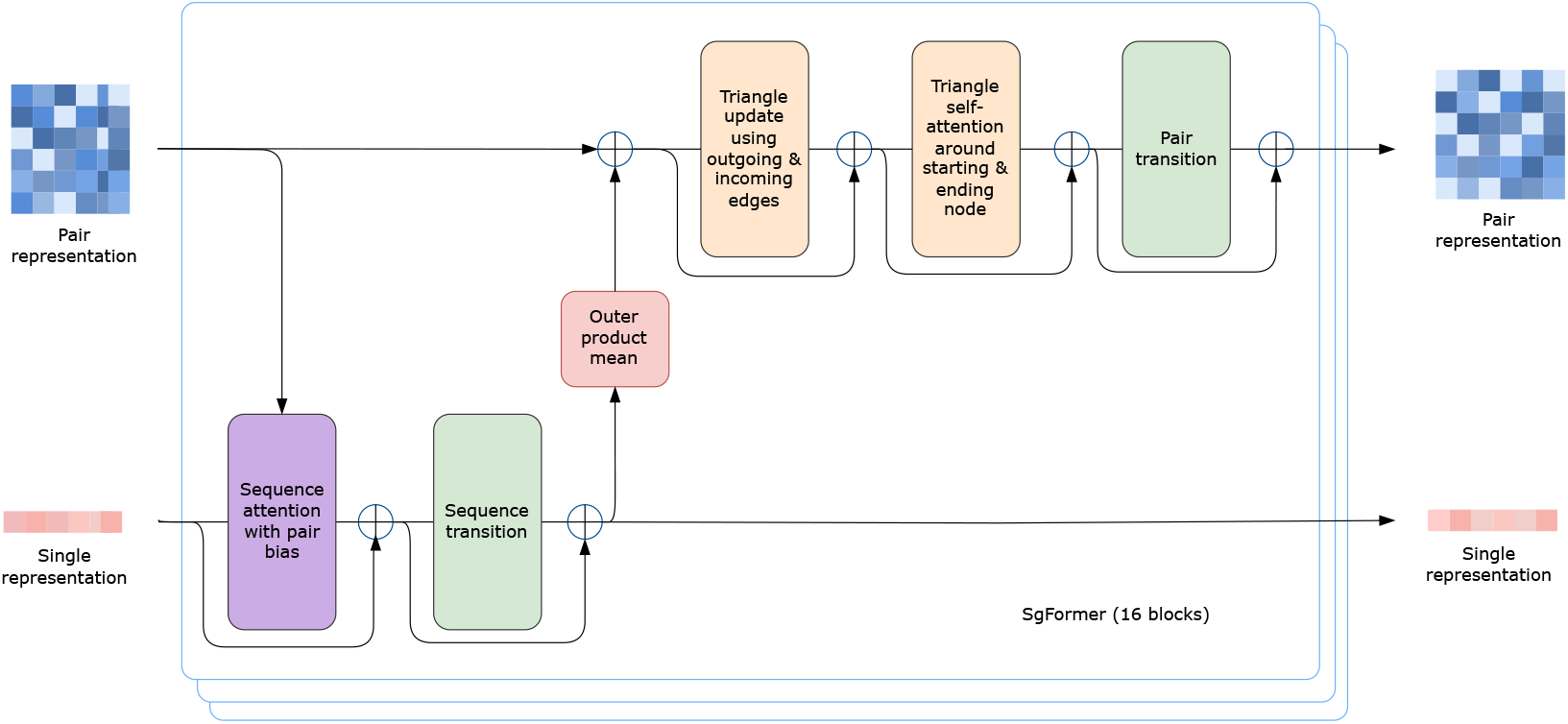
The SgFormer blocks update the single and pair representations through sequence attention and triangle update module.

**Fig. A7:**
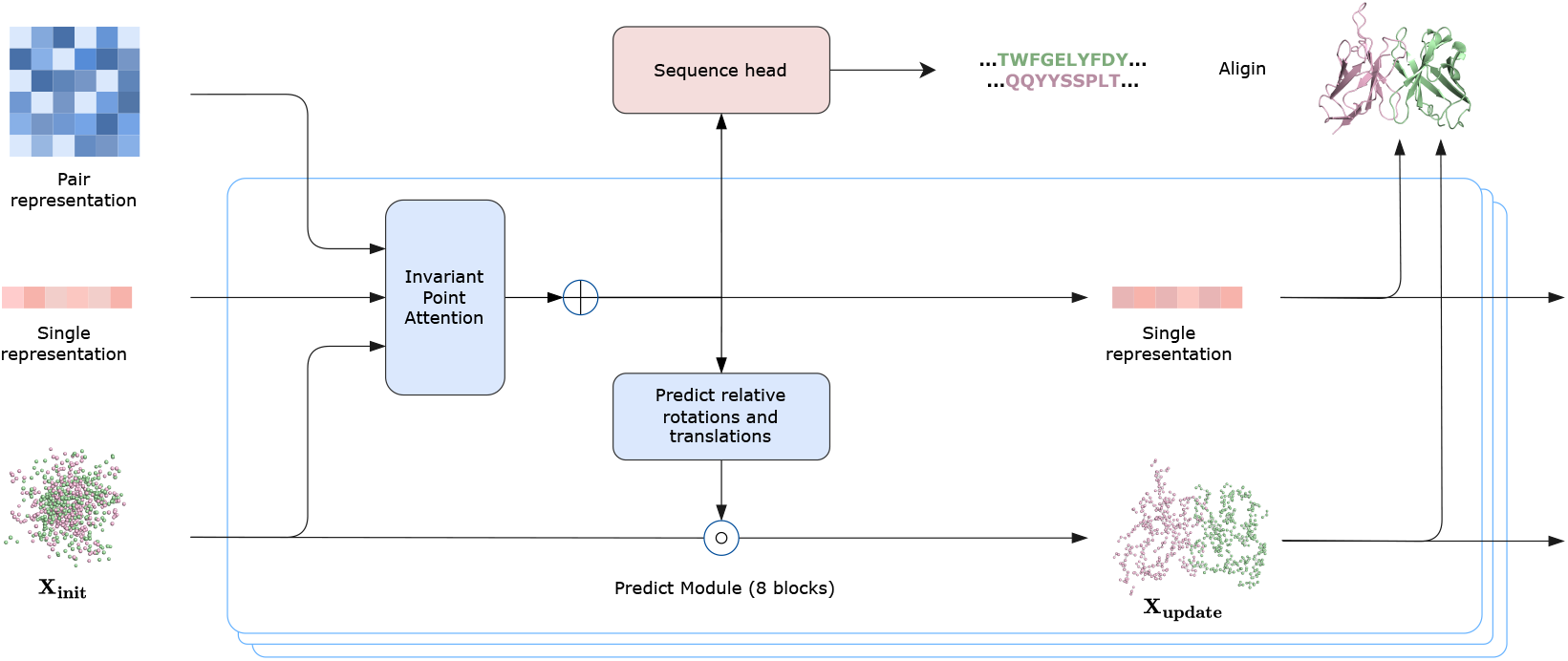
Antibody Sequence-Structure Prediction Module: The features input into a prediction module that includes an IPA module and a Sequence head to predict the sequence and structure of the antibody.

**Fig. A8:**
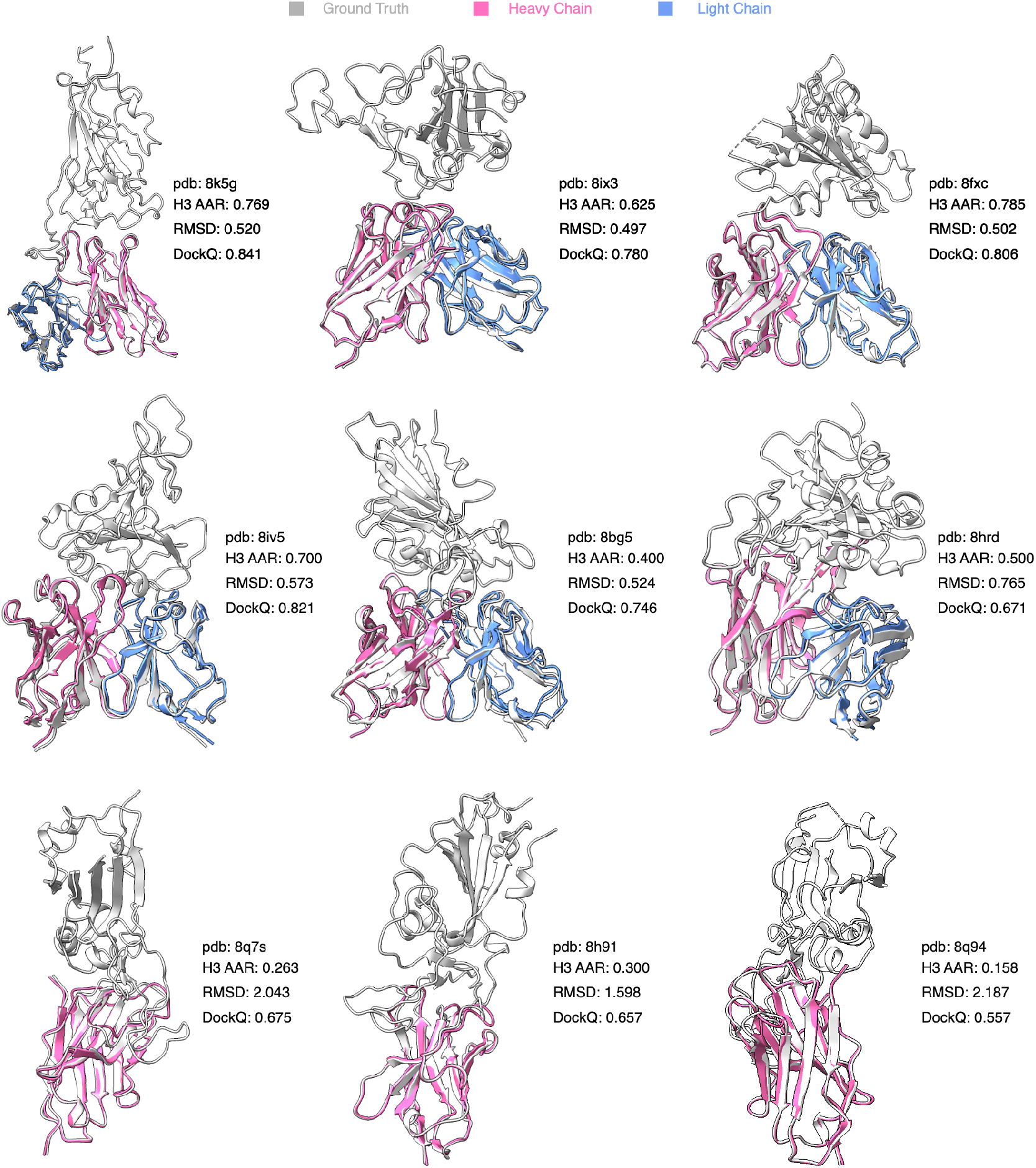
Samples of generated antibodies and nanobodies by IgGM.

## B Extended Data Tables

**Table B1:** List of antigen-antibody complexes in SAb-23H2-Ab.

|  |  |  |  |  |  |
| --- | --- | --- | --- | --- | --- |
| 8fxc_h.l.e | 8d4r_h.l.g | 8k5g_h.l.a | 8iv5_a.b.g | 8g4p_c.d.e | 8h7z.f.i.a |
| 8szy_c.d.u | 8j7y.f.d.b | 8iv4_a.b.g | 8wsq_a.b.f | 8g4t_h.l.i | 8dpl_o.o.e |
| 8fdo_a.b.c | 8hpu_m.n.a | 8dyy_e.f.i | 8iv8_h.l.g | 8g4p_a.b.e | 8t04_e.f.b |
| 8dz3_h.l.i | 8t9z_h.l.a | 7trh_h.b.g | 8ucd_h.l.b | 8ix3_h.l.g | 8bg5_a.a.b |
| 8r8d_e.i.b | 8gh4_h.l.e | 8szy_i.m.u | 8eay_h.l.a | 8hrd_n.q.k | 7yv1_h.l.a |
| 8gsp_h.l.2 | 8b7h_h.l.a | 8dpz_f.g.b | 8dyw_c.d.i | 7yh6_h.l.a | 8j7e_a.b.f |
| 8pq2_h.l.c | 8ezl_h.h.a | 8ded_u.v.f | 8eee_a.b.z | 8ez7_h.l.a | 8tco_f.g.c |
| 8d0y_h.l.g | 8udg_h.l.a | 8ezm_h.h.a | 8e6j_e.i.b | 8awm_h.l.a | 8dun_i.m.k |
| 8smi_h.l.a | 8byu_h.l.a | 8e6k_h.l.a | 8f5i_x.y.a | 8tea.f.i.d | 8hrd_w.x.t |
| 8tea_h.g.c | 8gye_e.f.a | 8ee5_h.l.z | 8grr_h.l.2 | 8x0t_h.l.a | 8fsj_c.b.e |

**Table B2:** List of antigen-antibody complex in SAb-23H2-Nano.

|  |  |  |  |  |  |
| --- | --- | --- | --- | --- | --- |
| 8q94_c.na.a | 8fcz_c.na.a | 7z1x_c.na.a | 8q7s_l.na.j | 8ee2_g.na.e | 8h91_d.na.b |
| 8q7s_e.na.d | 8h5t_b.na.a | 8aix_d.na.k | 8sk5_b.na.a | 7y0o_b.na.a | 8hr2_b.na.a |
| 8elq_b.na.a | 7zau_d.na.c | 8hbj_e.na.c | 7ymh_d.na.a | 8q95_b.na.a | 7z1x_b.na.a |
| 8hr2_c.na.a | 8eln_g.na.e | 8b18_b.na.a | 8g0w_d.na.b | 8h5u_d.na.c | 8h64_d.na.c |
| 8c9x_h.na.c | 8c5h_n.na.s | 8q93_c.na.a |  |  |  |

**Table B3:**
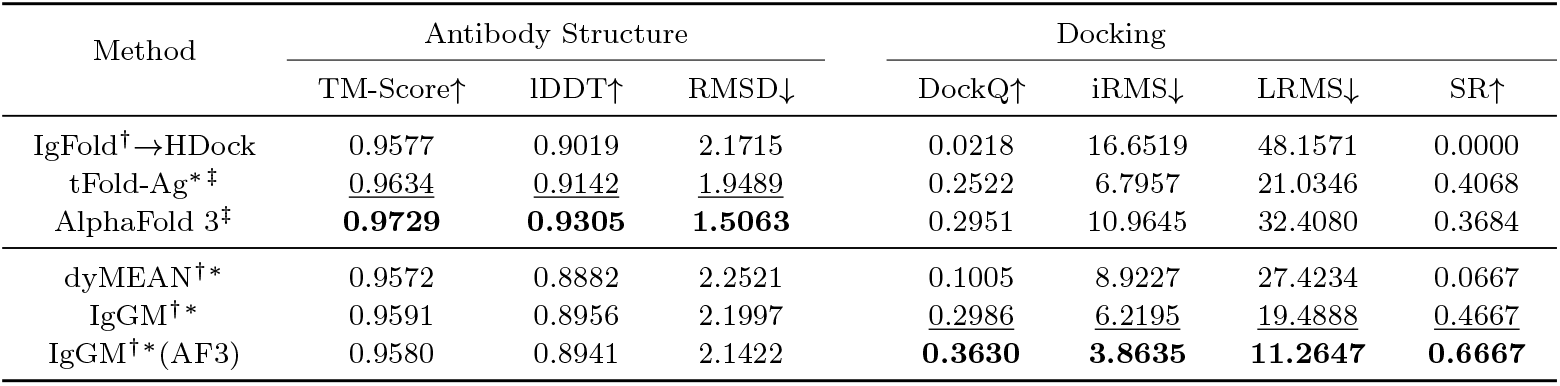
Complex structure prediction. Methods with superscript ^*†*^use antigen structure as input, while methods with superscript ^*∗*^ utilize epitope information as input; methods with superscript ^*‡*^ use multiple sequence alignment (MSA). (AF3) indicates that the structure predicted by AlphaFold 3 is used as the initial input. The root mean square deviation (RMSD) in CDR H3 is reported. **Bold** indicates the best performance, while *<u>underline</u>* indicates the second-best performance.

**Table B4:** Results of the novel antibody design on SAb-2023H2-Ab. (IgFold) or (AF3) indicates that the antibody structure predicted by IgFold or AlphaFold 3 is used as the initial input. Backbone RMSD in different CDR regions are reported. H1-H3 indicate the CDRs of heavy chain while L1-L3 indicate the CDRs of light chain. **Bold** indicates the best performance, while <u>underline</u> represents the second best.

| Model |  | DiffAb<br>(IgFold) | DiffAb<br>(AF3) | MEAN<br>(IgFold) | MEAN<br>(AF3) | dyMEAN | IgGM | IgGM<br>(AF3) |
| --- | --- | --- | --- | --- | --- | --- | --- | --- |
| AAR↑ | L1 | 0.597 | 0.608 | - | - | 0.633 | **0.750** | <u>0.737</u> |
|  | L2 | 0.598 | 0.599 | - | - | 0.634 | **0.743** | <u>0.735</u> |
|  | L3 | 0.421 | 0.424 | - | - | 0.570 | **0.635** | <u>0.602</u> |
|  | H1 | 0.642 | 0.637 | - | - | **0.742** | <u>0.740</u> | 0.739 |
|  | H2 | 0.363 | 0.394 | - | - | 0.627 | **0.644** | <u>0.639</u> |
|  | H3 | 0.214 | 0.226 | 0.248 | 0.246 | 0.294 | **0.360** | <u>0.330</u> |
| RMSD↓ | L1 | 0.783 | 0.749 | - | - | 0.864 | **0.589** | <u>0.659</u> |
|  | L2 | 0.471 | 0.466 | - | - | 0.481 | **0.378** | <u>0.395</u> |
|  | L3 | 1.002 | 1.017 | - | - | 0.941 | **0.847** | <u>0.903</u> |
|  | H1 | 0.650 | 0.623 | - | - | 0.633 | **0.555** | <u>0.590</u> |
|  | H2 | 0.641 | 0.586 | - | - | 0.705 | **0.486** | <u>0.566</u> |
|  | H3 | 2.741 | 2.646 | 2.357 | 2.300 | 2.454 | **2.131** | <u>2.155</u> |
| Docking | DockQ↑ | 0.022 | 0.208 | 0.022 | 0.207 | 0.079 | <u>0.246</u> | **0.326** |
|  | iRMS↓ | 17.034 | 9.731 | 16.838 | 8.968 | 9.698 | <u>6.579</u> | **4.030** |
|  | LRMS↓ | 48.163 | 27.559 | 48.104 | 27.557 | 28.764 | <u>19.678</u> | **11.229** |
|  | SR↑ | 0.000 | 0.368 | 0.000 | 0.354 | 0.049 | <u>0.433</u> | **0.627** |

**Table B5:** Ablations performance for IgGM.

| Method | AAR↑<br>CDR3 | DockQ↑ | iRMS↓ | LRMS↓ | SR↑ |
| --- | --- | --- | --- | --- | --- |
| IgGM | 0.360 | 0.246 | 6.579 | 19.678 | 0.433 |
| w/o two stage training | 0.160 | 0.072 | 10.260 | 30.961 | 0.000 |
| w/o epitope | 0.326 | 0.069 | 14.609 | 36.967 | 0.050 |
| w/o PPSM | 0.322 | 0.233 | 7.444 | 20.996 | 0.426 |
| w/o mixed training | 0.334 | 0.231 | 7.524 | 22.713 | 0.350 |

**Table B6:** Different sample steps for IgGM.

| Method | AAR↑<br>CDR3 | DockQ↑ | iRMS↓ | LRMS↓ | SR↑ |
| --- | --- | --- | --- | --- | --- |
| Step=1 | 0.362 | 0.240 | 6.474 | 19.694 | 0.383 |
| Step=2 | 0.363 | 0.244 | 6.570 | 19.616 | 0.383 |
| Step=5 | 0.363 | 0.242 | 6.610 | 19.932 | 0.400 |
| Step=10 | 0.360 | 0.246 | 6.579 | 19.678 | 0.433 |
| Step=20 | 0.361 | 0.232 | 6.840 | 20.980 | 0.400 |
| Step=50 | 0.348 | 0.225 | 6.809 | 20.905 | 0.383 |

**Table B7:** Results of structure prediction and novel nanobody design on SAb-2023H2-Nano. (AF3) signifies that the structure predicted by AlphaFold 3 is utilized as the initial input. Backbone RMSD in different CDR3. CDR1-CDR3 indicate AAR of the rigons. **Bold** indicates the best performance.

| Structure prediction of Nanobody |  |  |  |  |  |  |  |  |
| --- | --- | --- | --- | --- | --- | --- | --- | --- |
| Method | TM-Score↑ | lDDT↑ | RMSD↓ |  | DockQ↑ | iRMS↓ | LRMS↓ | SR↑ |
| tFold-Ag | 0.9344 | **0.9303** | 1.6722 |  | 0.2881 | **6.3490** | **15.0810** | 0.4296 |
| AlphaFold 3 | **0.9519** | 0.9286 | **1.1885** |  | 0.2867 | 11.2194 | 32.6760 | 0.3885 |
| IgGM | 0.9318 | 0.8931 | 1.9925 |  | **0.2907** | 7.9879 | 22.0168 | **0.4400** |
| Design of Nanobody |  |  |  |  |  |  |  |  |
| Method | CDR1↑ | CDR2↑ | CDR3↑ | RMSD↓ | DockQ↑ | iRMS↓ | LRMS↓ | SR↑ |
| DiffAb (AF3) | 0.533 | 0.291 | 0.156 | 2.274 | 0.211 | 13.265 | 35.805 | 0.346 |
| IgGM | **0.565** | **0.330** | **0.183** | **1.980** | **0.267** | **6.927** | **14.966** | **0.415** |

**Table B8:**
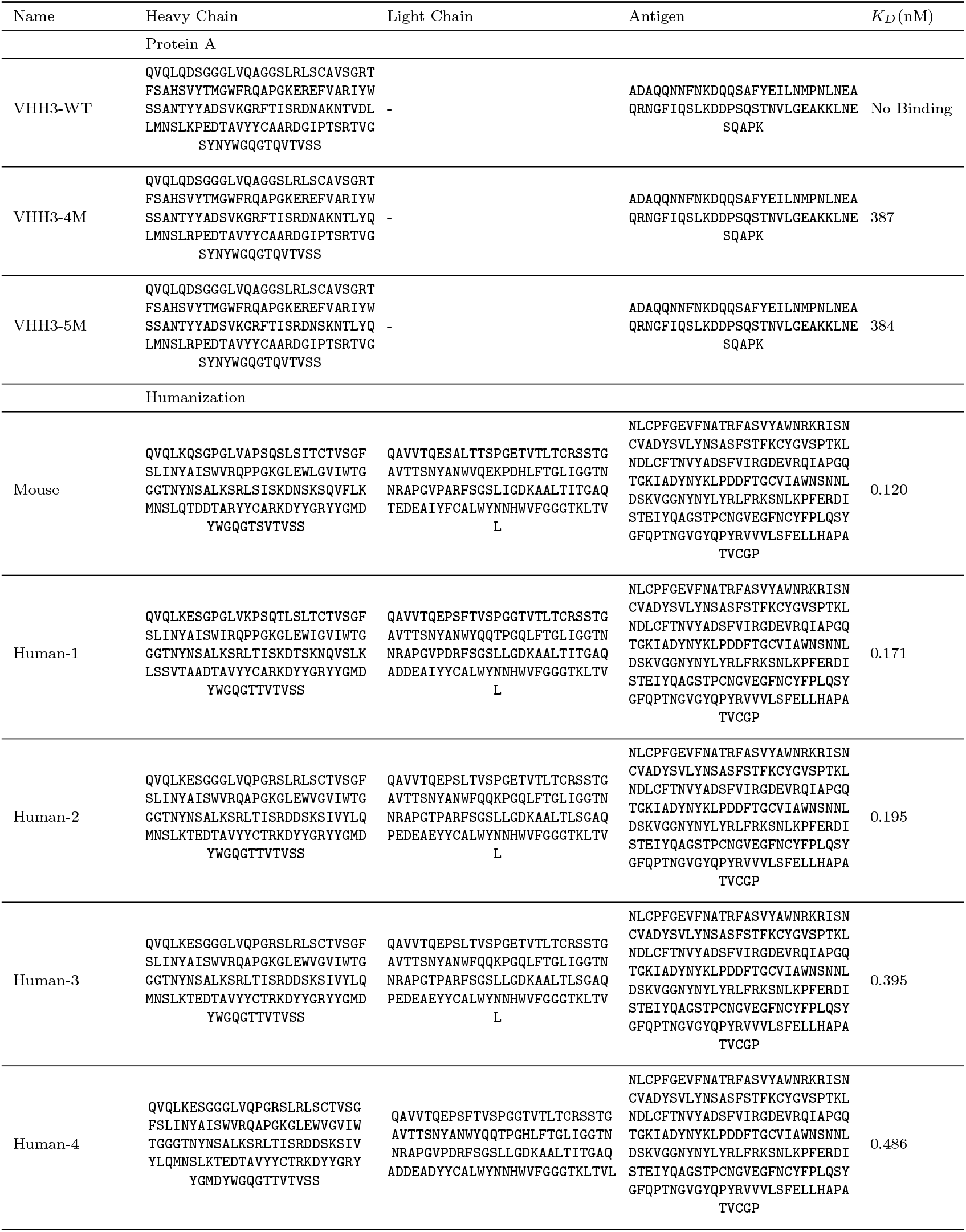
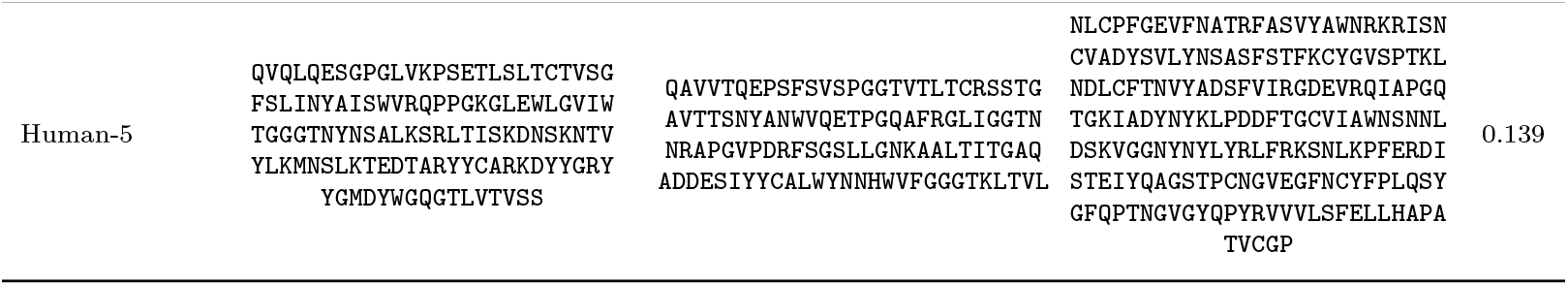
Wet experimental results of framework region design.

**Table B9:**
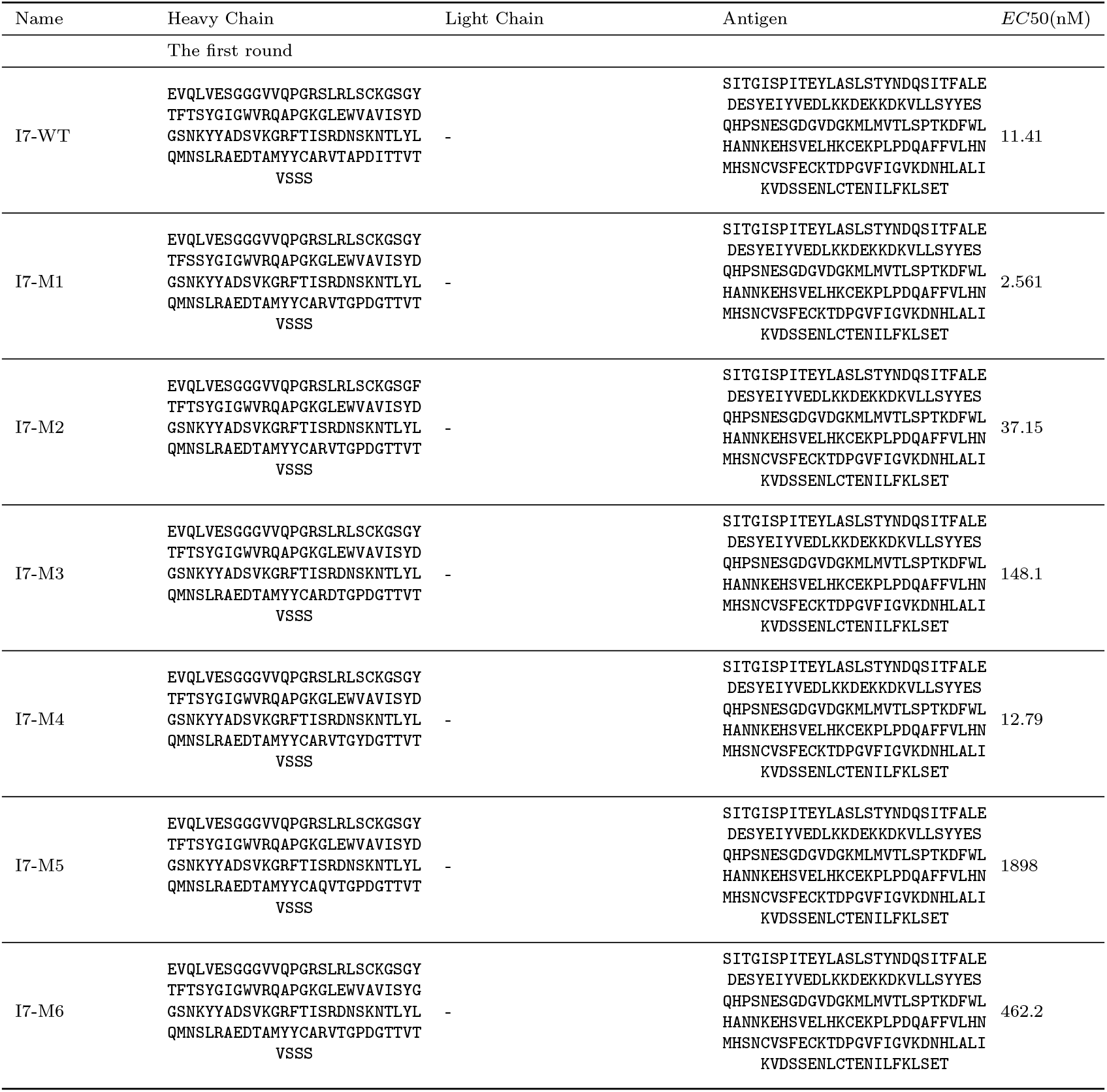
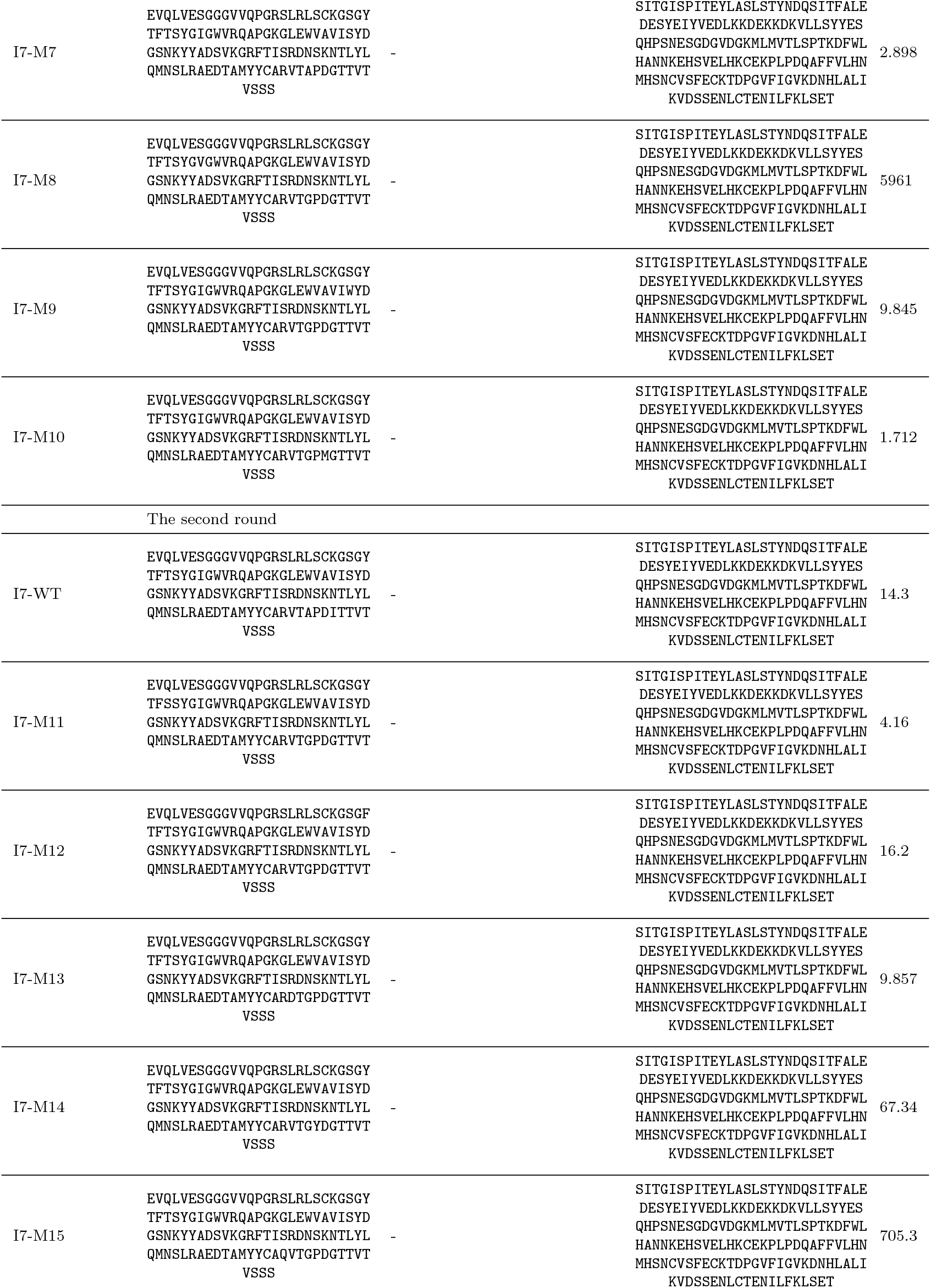
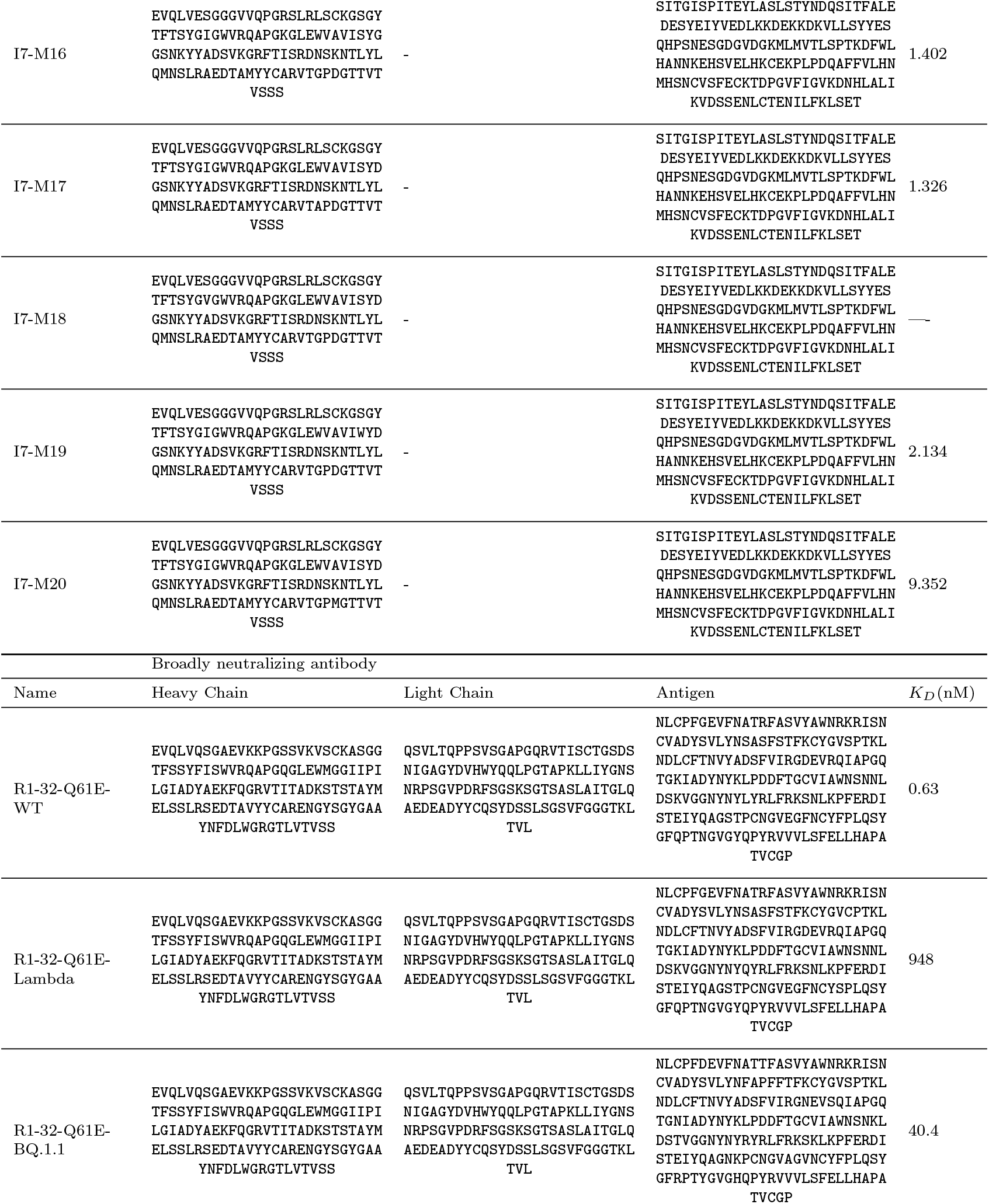
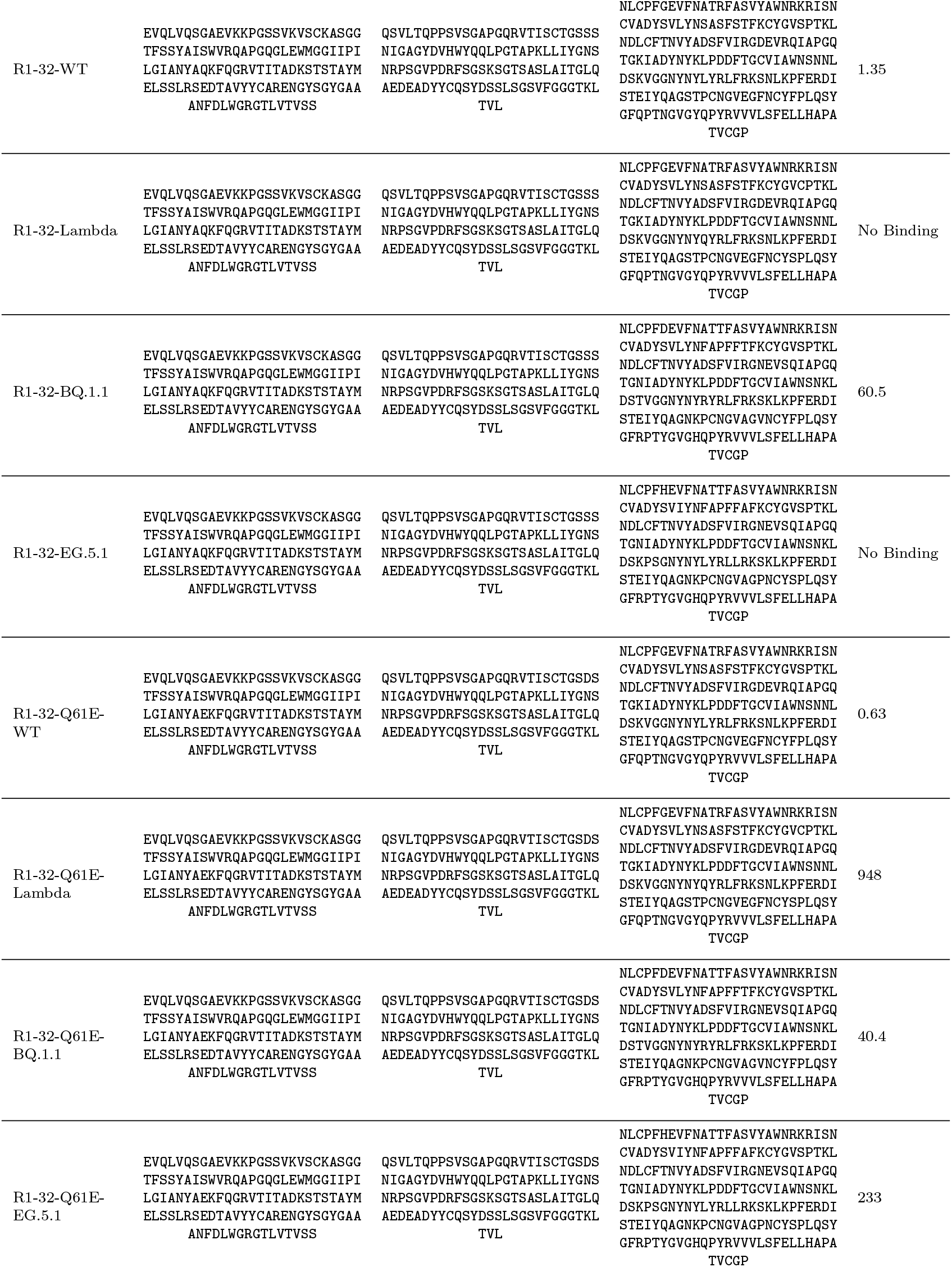
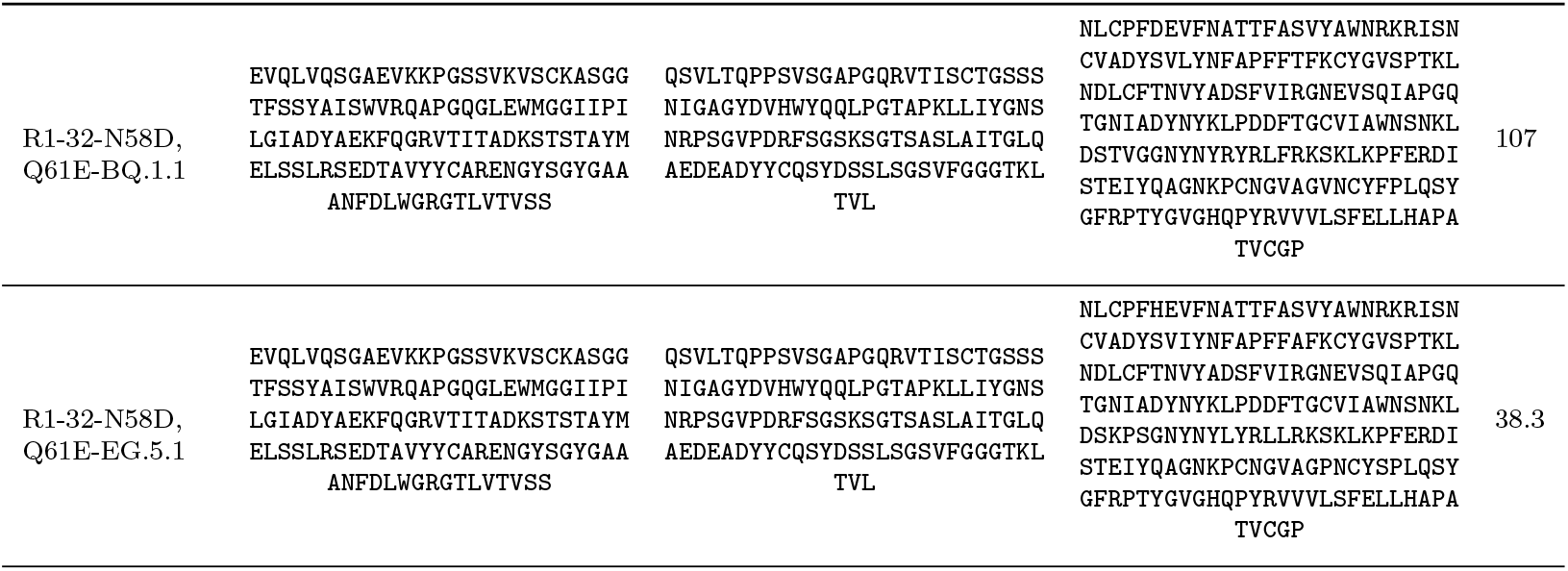
Wet experimental results of affinity maturation.

**Table B10:**
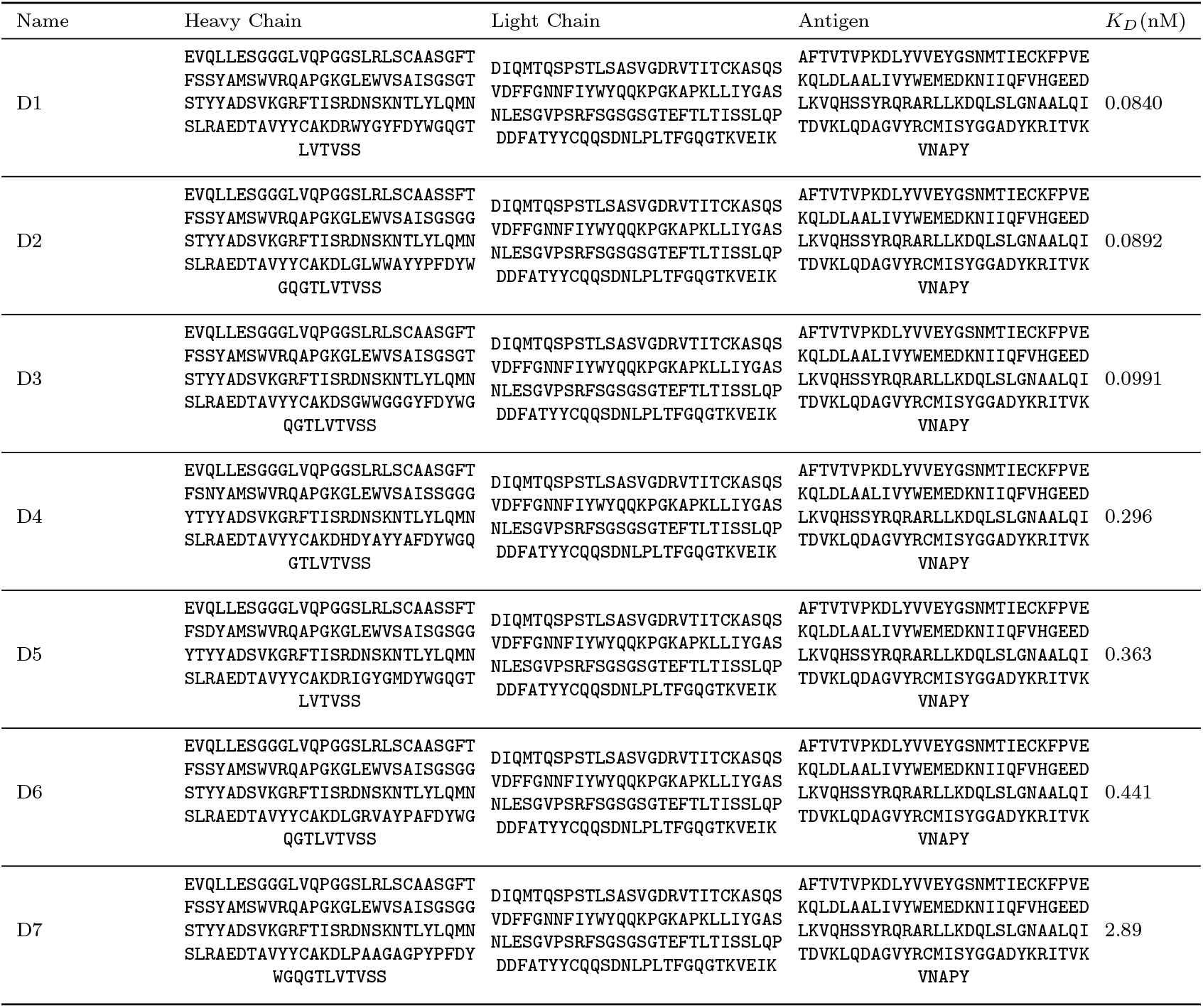
Wet experimental results of *de novo* design.

## C Supplementary Methods

### C.1 Details of testing comparative methods

In this section, we introduce the details of running the comparative methods.

**IgFold**. We use the official repository (https://github.com/Graylab/IgFold) of IgFold for testing.

**tFold**. We use the official repository (https://github.com/TencentAI4S/tfold) of tFold for testing.

**AlphaFold 3**. Due to the limitations of AlphaFold 3, we use the official server (https://alphafoldserver.com/) for testing.

**MEAN(IgFold)**. Since MEAN does not support input without antibody structures, we use the antibody structures predicted by IgFold [13] as the initial input for MEAN. We use the test code from the official GitHub repository (https://github.com/THUNLP-MT/MEAN) to generate the results.

**MEAN(AF3)**. Since MEAN does not support input without antibody structures, we use the antibody structures predicted by AlphaFold3 [11] as the initial input for MEAN. We use the test code from the official GitHub repository (https://github.com/THUNLP-MT/MEAN) to generate the results. It should be noted that thanks to the accuracy of the structures predicted by AlphaFold3, there is a significant improvement in the performance of the structural part metrics.

**DiffAb(IgFold)**. Since DiffAb does not support input without antibody structures, we use the antibody structures predicted by IgFold [13] as the initial input for MEAN. We use the test code from the official GitHub repository (https://github.com/luost26/diffab) to generate the results.

**DiffAb(AF3)**. Since DiffAb does not support input without antibody structures, we use the antibody structures predicted by AlphaFold3 [11] as the initial input for MEAN. We use the test code from the official GitHub repository (https://github.com/luost26/diffab) to generate the results. It should be noted that thanks to the accuracy of the structures predicted by AlphaFold3, there is a significant improvement in the performance of the structural part metrics.

**dyMEAN**. For dyMEAN, we conducted tests using the official GitHub repository (https://github.com/THUNLP-MT/dyMEAN). During the testing process, only the sequence of the antibody’s framework region was provided.

### C.2 Diffusion models

Diffusion models [50, 51, 68] are a type of generative model that have been successfully applied in various fields, including image generation [69–71], protein design [25, 72, 73], among others. Diffusion models gradually add noise to the data through a forward process until it becomes random noise, and then learn a reversible backward process to progressively recover the original data from the noise.

#### C.2.1 Continuous Diffusion

To model a distribution *p*(*w*), an effective method involves first embedding *w* into a continuous variable *x*_0_ using an embedding matrix *U*_*θ*_ and then adding Gaussian noise. For the *C*_*α*_ coordinates of antibodies, the values in three-dimensional space are continuous, and therefore a continuous diffusion model can be used to generate the *C*_*α*_ coordinates [49]. In practical applications, Gaussian noise can be gradually added to the coordinate values until they approach a Gaussian distribution. The prior distribution is set as *π*(*x*) = 𝒩 (0, *I*), and the forward process is defined by:

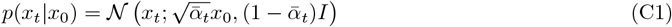

where 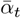 ranges from 0 to 1. The values of 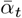 are determined by a predefined noise schedule, such as a linear or cosine schedule, which controls the rate at which noise is added over *T* time steps. For example, the noise schedule can be defined as 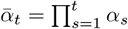, where *α*_*s*_ is the noise coefficient at step *s*. The forward process ensures that the distribution of *x*_*t*_ becomes increasingly Gaussian as *t* increases.

The reverse process aims to learn a function *p*_*θ*_(*ŵ*|*x*_*t*_, *t*) that can reconstruct the sequence from the noisy data points *x*_*t*_. This is achieved by minimizing the following loss function with respect to *θ*:

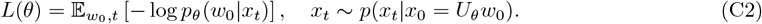

The loss function is typically approximated by training the model to predict the noise *ϵ* added during the forward process. Specifically, the model learns to minimize the mean squared error (MSE) between the predicted noise *ϵ*_*θ*_(*x*_*t*_, *t*) and the true noise *ϵ*:

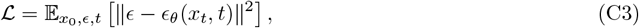

where 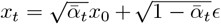. This formulation allows the model to iteratively denoise the data during the reverse process. Using *p*_*θ*_(*ŵ*|*x*_*t*_, *t*), we can define the reverse process distribution as follows:

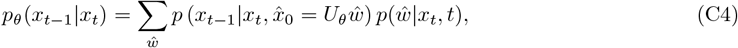

where *p*(*x*_*t−*1_|*x*_*t*_, *x*_0_) is also a Gaussian distribution. During inference, the learned reverse process can be used to transform samples from *π*(*x*) into samples from the learned distribution *p*_*θ*_(*x*_0_). This is done by iteratively sampling from *p*_*θ*_(*x*_*t−*1_|*x*_*t*_) and then sampling *w ∼ p*_*θ*_(*ŵ*|*x*_0_, 0).

#### C.2.2 Discrete Diffusion

To model a discrete sequence generation process for antibody sequences, we can leverage the framework of discrete diffusion models. As proposed by [53], for scalar discrete random variables with *K* categories, denoted as *s*_*t*_ and *s*_*t−*1_ where *s*_*t*_, *s*_*t−*1_ *∈ {*1, …, *K }*, the forward and reverse processes are defined using categorical distributions. For antibody sequences, there are 20 classes of amino acids, and each position in the sequence can be modeled as a categorical distribution over these 20 classes.

##### Forward Process

The forward process involves gradually adding noise to the sequence, transforming the initial sequence **s**_0_ into a noisy version **s**_*t*_. This is achieved by defining transition probabilities between states at consecutive time steps. For a specific antibody sequence, the amino acid type at each position can be regarded as a categorical distribution. For a random variable at a certain position, there are 20 classes, denoted as *s*_*t*_, …, *s*_*t−*1_ *∈ {*1, …, 20*}*.

The forward transition probabilities are represented using matrices:

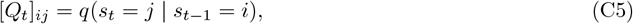

where *Q*_*t*_ is a *K × K* matrix encoding the probability of transitioning from state *i* to state *j* at step *t*. For antibody sequences, *K* = 20, corresponding to the 20 amino acid types.

Representing *s* in its one-hot form as a row vector **s**, the forward transition can be expressed as:

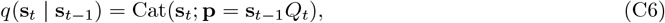

where Cat(**s**; **p**) denotes a categorical distribution over the one-hot vector **s** with probabilities from **p**. The term **s**_*t−*1_*Q*_*t*_ represents a row vector-matrix product, with *Q*_*t*_ applied independently to each position in the sequence. This formulation ensures that the noise is added in a way that preserves the categorical nature of the sequence.

##### Multi-Step Forward Transition

Starting from **s**_0_, the *t*-step marginal distribution is given by:

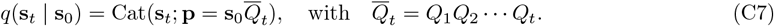

Here, 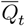 is the cumulative transition matrix after *t* steps, obtained by multiplying the individual transition matrices *Q*_1_, *Q*_2_, …, *Q*_*t*_. This ensures that the forward process is a Markov chain, where the state at step *t* depends only on the state at step *t −* 1.

##### Reverse Process

The reverse process aims to learn a function *p*_*θ*_(**s**_*t−*1_ | **s**_*t*_, **s**_0_) that reconstructs the original sequence from the noisy data points **s**_*t*_. This is achieved by minimizing the following loss function with respect to *θ*:

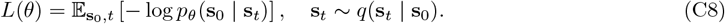

The loss function encourages the model to predict the noise added during the forward process, similar to the continuous diffusion case.

##### Posterior Distribution

The reverse process distribution is defined as:

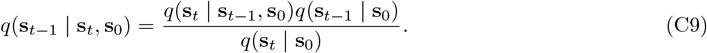

This posterior distribution can be simplified using the forward transition matrices:

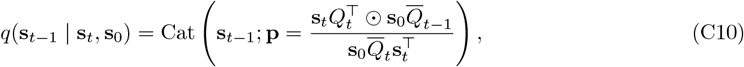

where ⊙ denotes element-wise multiplication (Hadamard product), and 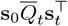 is a normalization factor ensuring the probabilities sum to 1.

#### C.2.3 SO(3) Diffusion

For each amino acid direction, the orientation within the local coordinate system of each amino acid can be considered as a continuous value in the SO(3) space, analogous to a uniform distribution on polar coordinates [52]. This representation leverages the fact that rotations in three-dimensional space form the compact Lie group SO(3), which is a non-Euclidean manifold. To model the diffusion of rotational data, we construct both forward and backward processes on this group, ensuring that the structural constraints (orthogonality and determinant conditions) are preserved throughout the process.

##### Forward Diffusion Process

The forward process diffuses the initial orientation data *o*_0_ *∈* SO(3) into pure noise over *T* time steps. At each step *t*, the distribution *q*(*o*_*t*_ | *o*_0_) is defined as an isotropic Gaussian distribution on SO(3) (denoted IGSO(3)), parameterized by a geodesic-based scalar product and a noise scale. Specifically:

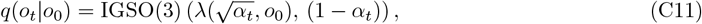

where: *α*_*t*_ *∈* [0, 1] is a time-dependent schedule that controls the noise intensity (*e*.*g*., *α*_*t*_ = exp(*−β*_*t*_) for some 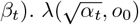 represents the scalar multiplication along the geodesic from the identity rotation *I ∈* SO(3) to *o*_0_, scaled by 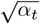. This ensures that the mean of the distribution evolves smoothly toward the identity as *α*_*t*_→ 0. 1*− α*_*t*_ determines the variance of the IGSO(3) noise, which increases as the process progresses. The isotropic Gaussian distribution on SO(3) (IGSO(3)) is constructed via the exponential map on the Lie algebra 𝔰𝔬(3). For a rotation matrix *R ∈* SO(3), the exponential map exp : 𝔰𝔬(3) → SO(3) generates a rotation by an angle *θ* around a unit vector *ω ∈* ℝ^3^, where *ω ∈* 𝔰𝔬(3) is represented as a skew-symmetric matrix. The IGSO(3) distribution samples a random skew-symmetric matrix Ω *∈* 𝔰𝔬(3) with Frobenius norm ∥Ω∥ _*F*_ = *σ*, and maps it to a rotation *R* = exp(Ω).

This approach ensures that the noise added at each step remains within the group SO(3), preserving the physical validity of orientations.

##### Backward Diffusion Process

The backward process aims to reverse the forward diffusion, transforming noisy orientations *o*_*T*_ back into meaningful data *o*_0_. At each step *t*, the distribution *p*(*o*_*t−*1_|*o*_*t*_, *o*_0_) is also modeled as an IGSO(3) distribution:

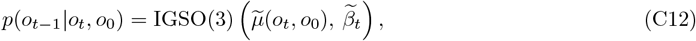

where 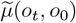 is the mean of the backward process, computed as the product of two rotation matrices:

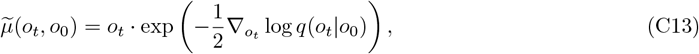

where 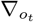 log *q*(*o*_*t*_|*o*_0_) is the gradient of the log-posterior with respect to *o*_*t*_, derived from the forward process. 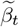 is the variance of the backward process, typically chosen to match the forward noise schedule *α*_*t*_, ensuring numerical stability.

The backward process is trained to approximate the true posterior *p*(*o*_*t−*1_|*o*_*t*_, *o*_0_) using a neural network. The network predicts the noise *ϵ*_*t*_ *∈* 𝔰𝔬 (3) added during the forward step, and the mean 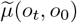 is computed as:

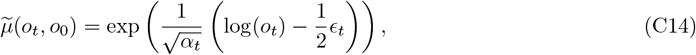

where log(*o*_*t*_) is the logarithmic map from SO(3) to 𝔰𝔬(3), and *ϵ*_*t*_ is the predicted noise vector.

### C.3 Consistency Models

Consistency Models (CMs) [67, 74] leverage the PF-ODE framework to create a direct relationship between data and noise distributions. The primary objective of CMs is to develop a consistency function *f* (**x**_*t*_, *t*) that effectively transforms a noisy image **x**_*t*_ back into its clean counterpart **x**_0_, adhering to a boundary condition at *t* = 0. This is accomplished through a specific parameterization:

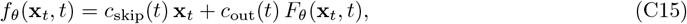

where the conditions *c*_skip_(0) = 1 and *c*_out_(0) = 0 ensure compliance with the boundary requirement. The functions *c*_skip_(*t*) and *c*_out_(*t*) are designed to satisfy these constraints while remaining differentiable with respect to *t*. This parameterization allows the model to retain the original signal **x**_*t*_ when *t* approaches 0 (*i*.*e*., *f*_*θ*_(**x**_0_, 0) = **x**_0_) and to gradually shift toward learning the mapping from noise to data as *t* increases. The core idea of CMs is rooted in the PF-ODE framework, which describes a smooth trajectory from the data distribution *p*_0_(*x*) to a tractable noise distribution *π*(*x*) (typically Gaussian). Specifically, the PF-ODE is defined as:

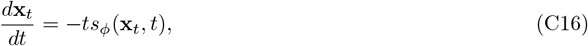

where *s*_*ϕ*_(**x**_*t*_, *t*) is the score function approximating *∇* log *p*_*t*_(**x**_*t*_). Unlike traditional diffusion models that rely on iterative sampling via SDEs, CMs directly model the inverse mapping from **x**_*T*_ (noisy input at time *T*) to **x**_0_ (clean output). This is achieved by training the consistency function *f*_*θ*_ to satisfy the self-consistency property:

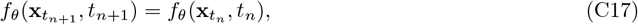

for any two points 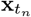 and 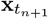 along the PF-ODE trajectory. This property ensures that the model’s predictions are invariant to the choice of time step along the same trajectory, enabling single-step generation or multi-step refinement.

During training, the PF-ODE is discretized into *N−* 1 segments, and a loss function is minimized to quantify the difference between adjacent points along the sampling path:

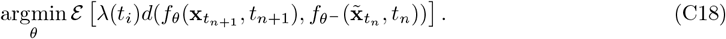

Here, *d*(*·, ·*) represents a chosen metric (*e*.*g*., L1, L2), *f*_*θ*_*−* denotes an exponential moving average (EMA) of previous outputs, and 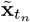 is generated via the PF-ODE forward process:

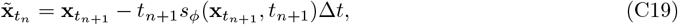

where Δ*t* = *t*_*n*+1_*− t*_*n*_. The use of EMA for *f*_*θ*_*−* is critical to prevent the model from collapsing to trivial solutions (*e*.*g*., mapping all inputs to zero), as demonstrated in the Consistency Distillation (CD) framework. In CD, the model learns to match its predictions with those of a teacher model derived from a pre-trained score-based diffusion model. This approach avoids the need for large-scale pretraining and reduces computational costs.

For further optimization, the Consistency Training (CT) strategy is introduced, which eliminates the dependency on a teacher model by directly using the PF-ODE forward process to generate 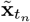. This method simplifies the training pipeline while maintaining the self-consistency property. The loss function in CT is modified to:

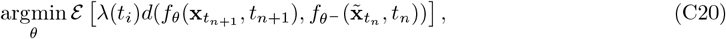

where 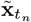 is now derived purely from the PF-ODE forward dynamics without external supervision. A key advantage of CMs is their ability to perform zero-shot image editing tasks, such as inpainting, colorization, and super-resolution, without task-specific training. For example, in image inpainting, a mask matrix *M* is applied to the noisy input **x**_*T*_, and the model iteratively refines the masked region by enforcing consistency across time steps. Similarly, for super-resolution, a low-resolution input is upsampled and treated as a noisy version of the high-resolution target, with the model learning to restore fine details through multi-step sampling. The sampling process in CMs can be executed in two modes: single-step generation and multi-step refinement. In single-step generation, the model directly maps **x**_*T*_ *∼* 𝒩 (0, *T* ^2^*I*) to **x**_0_ via:

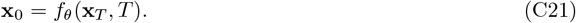

However, this approach may sacrifice detail quality due to the abrupt transition from noise to data. To address this, multi-step refinement is employed, where the model alternates between prediction and noise addition (via the PF-ODE forward process) to progressively refine the output. For a sequence of time steps *τ*_1_ *> τ*_2_ *> · · · > τ*_*N*_, the process is defined as:

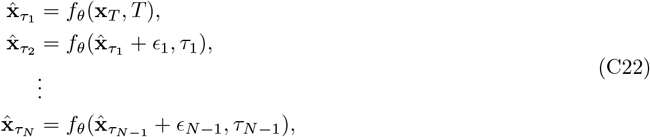

where 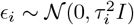 introduces controlled noise at each step. This iterative refinement balances speed and quality, making CMs highly versatile for applications requiring both efficiency and fidelity.

## D Experiment details

### D.1 Dataset

We selected all experimentally determined antibody structures published in the SAbDab database [29] up to December 31, 2022, as our training set. The final training set consisted of 6,448 antibody-antigen complexes with both heavy and light chains and 1,907 single-chain antibody-antigen complexes, ensuring a balanced representation of canonical and simplified antibody architectures. During the training process, we employed CD-Hit [48] to cluster the training set based on sequence similarity. Specifically, we set a clustering threshold of 95% sequence identity, grouping antibodies with high homology into clusters to mitigate overfitting risks. This process yielded 2,436 clusters, each containing structurally and functionally related antibody variants. To ensure efficient utilization of the data while maintaining diversity, we randomly sampled one representative sample per cluster for training in each epoch, thereby avoiding redundancy while preserving the heterogeneity of the dataset.

The validation and test sets were carefully curated to reflect the most recent experimental data. The validation set included antigen-antibody complexes determined experimentally and published between January 1, 2023, and June 30, 2023, while the test set comprised data from July 1, 2023, to December 30, 2023. To eliminate redundancy and ensure an unbiased evaluation, we removed sequences with sequence identities exceeding 90% to those in the training set using a custom filtering pipeline. This rigorous curation resulted in 101 validation samples and 60 test samples, both of which were completely disjoint from the training set in terms of sequence and structural overlap. The validation set was explicitly used for hyperparameter tuning and model selection, while the test set, named SAb-23H2-Ab, served as the benchmark for final performance evaluation.

To further validate the generalizability of our approach, we also constructed a specialized test set for nanobodies using a dataset released in the second half of 2023, referred to as SAb-23H2-Nano. This dataset leveraged the unique structural properties of nanobodies, such as their compact size and extended CDR3 regions (16–18 amino acids), which enable access to cryptic epitopes inaccessible to conventional antibodies. The inclusion of nanobodies in the test set allowed us to evaluate the model’s ability to generalize across diverse antibody formats, particularly those derived from camelid immunization protocols. For detailed methodologies on data preprocessing, clustering parameters, and quality control measures, refer to Section D.4. This alignment strategy facilitated precise comparison of antigen-binding regions, particularly the complementarity-determining regions (CDRs), critical for antigen recognition.

### D.2 Model details

#### D.2.1 Pre-trained Protein Sequence Model

For the further pre-training of PPSM, a combination of four distinct datasets is utilized:

- **UniRef50** [75]: The dataset employed is the March 2023 version of UniRef50. It contains roughly 60 million monomeric sequences. To guarantee the integrity and validity of our validation set, we randomly chose 4,000 sequences that had not been part of the training and validation sets of ESM2. These sequences were acquired by determining the difference set between UniRef50 and the UniRef100 dataset released in September 2021. Subsequently, these selected sequences were used to build both the validation and test sets.
- **PDB**: We used the January 2023 version of the Protein Data Bank (PDB). In total, it contains 188,000 sequences of solved protein multimers. From the PDB, we picked 4,000 pairs of interacting chains. These pairs were sourced from 4,000 complex structures that were released after 2022. The selected pairs were then used to form the validation and test set.
- **PPI**: This dataset aggregates 1.3 million pairs of protein sequences that are known to potentially interact. The data is collected from a variety of existing databases. We combined pairs of interacting proteins from multiple sources including HINT [76], intACT [77], HIPPIE [78], prePPI [79], BioGRID [80], comPPI [81] and huMAP [82]. Since the data originates from diverse sources, redundancy is an inherent issue. To tackle this, we utilized MMSeqs2 (Steinegger et al., 2017) to eliminate duplicated PPI pairs based on 100% sequence identities. Additionally, to enhance data quality, we filtered out interactions with low confidence scores, specifically those with scores lower than 0.3 in intACT and comPPI, and those with a quality lower than 0.63 in HIPPIE. After these filtering steps, we selected 4,000 pairs for use in the validation and test phases.
- **Antibody**: This dataset consists of 1.5 million paired antibody sequences collected from the OAS [83]. However, considering that the CDR3 region of the antibody can be influenced by the antigen, and the OAS data fails to account for this factor, we did not include any of the paired antibody sequences in the validation or test set.

In the end, a total of 12,000 monomeric sequences along with their interaction pairs were acquired, and these were earmarked for the purposes of validation and testing. They were divided evenly into two parts. Specifically, half of them were chosen to form the validation set, while the other half was allocated to the test set. Such a balanced division plays a crucial role in ensuring that the performance of our model can be comprehensively and rigorously evaluated. To mitigate overfitting to some extent and prevent data leakage, we took measures to reduce the redundancy within the training data. Regarding the monomeric sequences in UniRef50, we eliminated certain sequences from the training set by following the procedure detailed in the reference [84]. The MMseqs2 search operation (with parameters *-min-seq-id 0*.*5 -alignment-mode 3 -max-seqs 1000 -s 7 -c 0*.*8 -cov-mode 0*) was carried out. In this operation, the training set served as the query database, and the validation and test set (which included monomeric sequences as well as each individual chain from multimers) was set as the target database. Any training sequence that had a 50% sequence identity match with a sequence in the validation and test set under this search condition was removed from the training set. As for the multimeric sequences sourced from Protein-Protein Interaction (PPI) data and the Protein Data Bank (PDB), we computed the length-weighted sequence similarity using the MMseqs2 search (with parameters *<u>-min-seq-id 0.9 -alignment-mode 3 -max-seqs 1000 -s 7 -c 0.8 -cov-mode 0</u>*). Given the relatively small quantity of PPI and PDB data, we opted for a more stringent 90% identity threshold when dealing with multimers.

To make the training of PPSM more efficient when dealing with a large quantity of residues in multimeric sequences, which poses challenges to GPU memory usage and computational efficiency, we utilized the *ContiguousCropping* algorithm borrowed from AlphaFold-Multimer [12]. In particular, for multimeric structures sourced from the PDB, we randomly pick two interacting chains and crop them proportionally based on their lengths, making sure that the structural contacts are maintained. When there is no structural data available for PPI sequence pairs, the cropping operation is only determined by the length of the sequences. For homomers, we use a consistent cropping approach across all chains to ensure uniformity. As for antibody sequence pairs, since they are usually shorter, the cropping step is not necessary. For monomeric sequences and PPI pairs lacking structural data, we apply a masking strategy similar to that used in conventional masked language models. During the training process, 15% of the amino acids are randomly selected. Among these selected amino acids, 80% are masked, 10% are replaced with a random residue, and the remaining 10% are left unchanged. To prevent data leakage in homomers, we ensure that the masking is consistent across all chains. When handling protein sequences that have experimental structures, we raise the probability to 30% that the residues at the contact interface between two chains are masked. This method enhances the model’s ability to learn about protein interactions from sequence data. For antibodies, considering the predictability of amino acid types in the framework regions, we adopt a strategy where residues in the CDRs loop have a 30% higher probability of being selected for masking, following the approach used in specialized antibody language models [85], while the framework regions stay the same. In the subsequent pre-training stage, we carried out a total of 128,000 training steps. Each step consists of a batch of 128 samples, which include individual sequences from UniRef50, pairs from multimeric PDB structures, PPI interaction pairs, and paired antibody sequences. The selection probabilities for these different types of samples are in the ratio of 2:3:3:2. We used the AdamW [86] optimizer with hyperparameters set as *β*_1_ = 0.9 and *β*_2_ = 0.999. The learning rate schedule starts with a linear warm-up period, increasing from 3e*−*6 to 3e*−*5 over the first 12,800 steps. After that, the learning rate remains fixed at 3e*−*5 for the rest of the training. To effectively manage the model across multiple GPUs, we implemented Fully Sharded Data Parallelism (FSDP) [87], which distributes both the model weights and the optimization states. This allowed us to increase the sequence crop size to 1024. For the validation and testing phases, we applied an Exponential Moving Average (EMA) [88] to the model parameters with a decay rate of 0.995, which helps in achieving smoother parameter updates. The model is selected based on achieving the lowest average loss on a combined validation subset. The entire subsequent pre-training was performed on a cluster of 8 NVIDIA A100 GPUs and was completed in approximately 30 hours. This setup enabled us to scale up the training process efficiently while making the most of the computational resources.

#### D.2.2 Inter-chain Feature Embedding Module

The input fed into the Inter-chain Feature Embedding Module consists of several key components. Firstly, there is a list of chain information denoted as chn infos, which likely contains various details about the chains relevant to the overall process. Secondly, an asymmetric ID vector, named asym id, is included. This vector presumably holds unique identifiers or characteristics related to the asymmetry in the system. Thirdly, the number of embedding dimensions, represented as *n dims*, is an important parameter that determines the dimensionality of the resulting feature tensor. And finally, the maximum relative index, ridx max, is also part of the input. The algorithm kicks off with the initialization of a linear layer. The input dimension of this linear layer is set to 2*×* ridx_max + 3, while the output dimension is set to *n dims*. This initial setup of the linear layer is crucial as it will play a significant role in transforming the incoming data. Following that, an asymmetric matrix is computed. This computation is based on the asym id. Once computed, the matrix is then converted into a floating-point tensor. This conversion is likely to make the data in a format that can be more easily processed in the subsequent steps. After that, an index vector is generated. This vector is used to calculate the relative index matrix. However, the calculated matrix might need some refinement. Thus, it undergoes a process of trimming and adjustment. Through these operations, the final relative index matrix is obtained. This matrix is then one-hot encoded, which is a common encoding technique to represent categorical data in a more suitable format for neural network processing. Subsequently, the asymmetric tensor (derived from the earlier steps) and the one-hot encoded tensor are concatenated along the last dimension. This concatenation combines different types of information into a single tensor. The combined tensor is then processed through the previously initialized linear layer. As a result of this processing, an updated feature tensor is produced. Finally, the algorithm outputs this updated tensor. This output tensor is then ready for subsequent feature processing. This entire process is highly effective as it manages to integrate the relative positions of the chains (through the relative index matrix) and the asymmetric information (from the asym id), thereby providing the model with rich and diverse contextual information that can potentially enhance the model’s performance in various tasks.

#### D.2.3 Single Channel Evoformer Module

The Sgformer module, which is a variant of the Evoformer, serves as the initial processing stage. In this stage, the Sgformer modules receive the feature provided by the PLM. These modules are carefully designed with a multi-layered architecture. Their purpose is to distill the essence of the antibody sequence. Through this architecture, they can effectively facilitate the learning of sequence features and pair-wise interactions. The sequence features that are extracted at this particular stage play a pivotal role. They are crucial for the subsequent task of recovering the original sequence from the perturbed input. On the other hand, the pair-wise representations are equally important. They encode the relational information, which is essential for understanding the complex folding of the antibody-antigen complex. This relational information helps in deciphering how the antibody and antigen interact and fold in relation to each other, which is a key aspect in understanding the biological mechanisms at play.

#### D.2.4 Structure Encoder

The structural module is composed of two distinct yet complementary components: one dedicated to encoding spatial information and the other focused on encoding interaction information between antigens and antibodies based on epitope information. For spatial encoding, the three-dimensional coordinates of each *C*_*α*_ atom across the protein structure are first extracted from the input structural data. These coordinates are then used to compute pairwise Euclidean distances between all *C*_*α*_ atoms within each sample, which results in the formation of a distance matrix *D ∈* ℝ^*N×N*^, where *N* denotes the number of residues in the structure. This matrix is subsequently transformed into a binary distance mask *M ∈ {*0, 1*}* ^*N×N*^ by applying a predefined threshold *τ* (*e*.*g*., *τ* = 8 Å), which distinguishes between proximal (contacting) and distal (non-contacting) residue pairs in the folded protein structure. The binary mask *M* serves as a critical structural prior that explicitly encodes spatial adjacency relationships, enabling the model to prioritize interactions between residues within the defined cutoff range. In addition to this binary representation, the continuous distance values within the distance tensor *D* are further discretized into *K* equidistant bins using a piecewise constant mapping function *f*_*d*_ : ℝ → *{*0, 1, …, *K −* 1*}*, where *K* is determined based on the empirical distribution of distances in the training dataset. Each distance value *d*_*ij*_ *∈ D* is mapped to its corresponding bin index 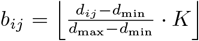, effectively converting the continuous distance metric into a categorical variable that preserves relative proximity information. These bin indices are then passed through an embedding layer 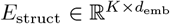, where *d*_emb_ represents the dimensionality of the learned structural embeddings, to generate a learnable representation *E*_*ij*_ = *E*_struct_(*b*_*ij*_) for each residue pair. The resulting structural encoding 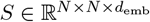 is further combined with the binary distance mask *M* through element-wise multiplication, ensuring that only interactions within the predefined spatial threshold contribute to the final representation. This dual-stage processing of spatial information—combining both binary proximity constraints and learnable distance-based embeddings—enables the model to simultaneously capture global structural context and fine-grained geometric relationships.

Epitope information encoding is utilized to generate embedded representations of the interaction and contact map features between antigens and antibodies. Initially, the input epitope information, which typically includes labeled residue positions on the antigen surface predicted to be involved in antibody binding, is mapped to a specified feature dimension *d*_in_ through a linear projection layer 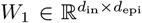, where *d*_epi_ denotes the original dimensionality of the epitope features (*e*.*g*., binary labels or physicochemical property vectors). This projection step ensures compatibility between the input epitope data and the subsequent processing layers. The mapped features are then further projected to the target dimension *d*_target_ using a second linear transformation 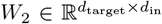, which aligns the feature space with the model’s hidden dimensions. During the transmission of epitope information, the module preprocesses the input epitope data to generate the corresponding feature representations through a domain-specific normalization strategy. Specifically, an appropriate feature generation method is selected based on the dimensionality of the input features: for high-dimensional epitope features (*e*.*g*., *d*_epi_ *>* 32), principal component analysis (PCA) is applied to reduce redundancy and retain the most informative components; for low-dimensional features (*e*.*g*., *d*_epi_ ≤ 8), feature expansion techniques such as polynomial basis functions are employed to enhance expressiveness. This preprocessing ensures that the type of features (*e*.*g*., sparse vs. dense) is consistent with the weight initialization strategy of the projection layer, preventing numerical instability during training. The preprocessed features are then passed through a hierarchical embedding layer 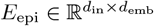 followed by a learnable projection layer 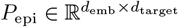, which jointly transforms the input into a continuous latent representation *Z* = *P*_epi_(*E*_epi_(*X*_epi_)), where *X*_epi_ denotes the preprocessed epitope data. The final feature embedding *Z* integrates both the discrete labeling of epitope residues and their contextual relationships with neighboring residues, enabling the model to discern subtle patterns in epitope topology. Through these steps, the epitope information encoding module effectively processes and transforms the input epitope data into a structured representation that captures both local binding preferences and global antigen-antibody interaction motifs. This enriched feature set not only facilitates the precise localization of epitopes but also guides the generative model in synthesizing antibodies with optimized binding affinity and specificity by aligning the encoded epitope features with the spatial constraints derived from the structural module.

### D.3 The objective of IgGM

For amino acid sequences, given that there are only 20 types of amino acids, different amino acids exhibit similar backbone atoms. We treat sequence recovery as a classification problem. We employ cross-entropy loss to guide the model in learning the correct sequence.

When it comes to the structure of antibodies, as previously mentioned in Section C.2, our objective is to recover the spatial coordinates of alpha carbons and the orientations of backbone atoms. To this end, we utilize the residue Frame Mean Squared Error (FMSE) loss, which has been demonstrated to be effective in [49]. This loss function is specifically designed to measure the discrepancy between the predicted and actual frames of protein residues, which are essential for accurately modeling the three-dimensional structure of antibodies. We have further developed an enhanced loss function termed inter-chain Frame Mean Squared Error (iFMSE) loss. This loss function is specifically tailored to impose constraints on the differences between different chains within the antibody structure. The iFMSE loss is designed to ensure that the model accurately captures the relative orientations and positions of the various chains that make up the antibody, which is crucial for maintaining the integrity of the quaternary structure. Additionally, we employ the inter-residue distance and angle metrics to reconstruct the orientations of backbone atoms. By incorporating these loss functions, our model is trained to not only accurately predict the amino acid sequences but also to reconstruct the intricate three-dimensional structures of antibodies, thereby enhancing the potential of our approach in the fields of structural biology and antibody modeling.

The overall objective of the model is to recover the correct sequence and structure from the denoised antibody data. The overall loss function can be represented as follows:

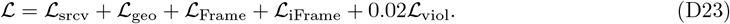

- **Amino-acid Sequence Recovery Loss** ℒ_**srcv**_ is designed to supervise the model to recover the amino-acid sequence *s*_*i*_ at the position *i*. A total 20 classes for common amino acid types are considered. Sequence embedding *{s*_*i*_*}* are linearly projected into the output classes and scored with the cross-entropy loss:

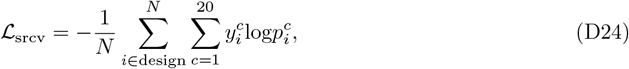

where 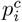 are predicted class probabilities, 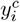 are one-hot encoded ground-truth values, and averaging across the masked positions.
- **Inter-residue Geometric Loss ℒ**_**geo**_ is designed to provide more direct supervision in the following stack. Four auxiliary heads, implemented as feed-forward layers, are added to the top of the final pair features for predicting inter-residue distances and angles, as described in trRosetta [89]. These include:
  – *D*_*ij*_: Distance between *C*_*β*_ and 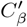
  – Ω_*ij*_: Dihedral angle formed by 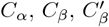, and 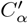
  – *Θ*_*ij*_: Dihedral angle formed by *N, C*_*α*_, *C*_*β*_, and 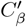
  – *Φ*_*ij*_: Planar angle formed by *C*_*α*_, *C*_*β*_, and 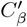 Each prediction head outputs a probabilistic estimation of the aforementioned distance and angles, denoted as 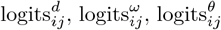, and 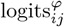. The cross-entropy loss is calculated for each term and summed up to form the final inter-residue geometric loss:

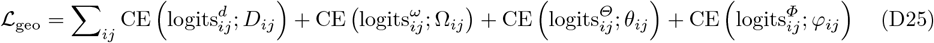
- **Residue Frame MSE Loss** is designed to provide more direct supervision in the predict module to recover the structure of antibody. The formula can be expressed as follows:

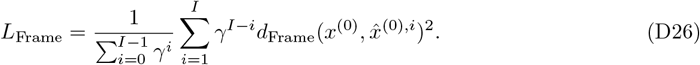

where 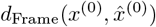 represents the distance about both rotation and translation, and it can be expressed in the following form:

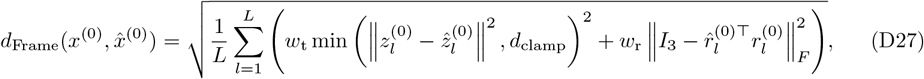

where *w*_t_ and *w*_r_ are weights on the translation and rotation distances, and *d*_clamp_ is a maximum distance to avoid the numerical overflow.
- **Interface Residue Frame MSE Loss** is designed to provide more direct supervision in the predict module to recover the structure of antibody between interchain. The formula can be expressed as follows:

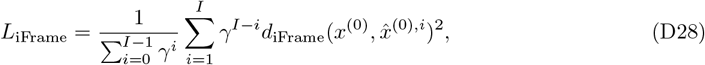

where 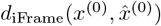 represents the distance about both rotation and translation, and it can be expressed in the following form:

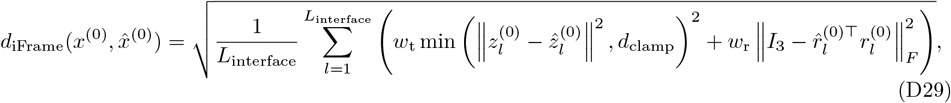

where *w*_t_ and *w*_r_ are weights on the translation and rotation distances, and *d*_clamp_ is a maximum distance to avoid the numerical overflow, *L*_interface_ represents the amino acid sequence of the contact region.
- **Structure violation loss** ℒ_**viol**_: Similar to AlphaFold2 [45], we introduce penalty terms for incorrect peptide bond length and angles, as well as steric clashes between non-bonded atoms. For multimer structure prediction, we do not penalize the bond length and angle between the last residue in the heavy chain and the first residue in the light chain, since there is no peptide bond between them. Besides, we normalize the steric clash loss by the number of non-bonded atom pairs in clash to stabilize the model optimization, as suggested in AlphaFold-Multimer [12].

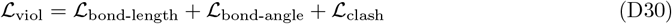

### D.4 Training Details

The training process of IgGM is divided into two stages, following the concept of curriculum learning. Initially, we train the model for structural design, using the Adam [86] optimizer with a default learning rate of *η* = 0.001, which is empirically validated to balance convergence speed and stability. The batch size is set to 32 to ensure sufficient gradient diversity while maintaining computational efficiency on 8 A100 GPUs. To enhance parameter robustness, we employ an Exponential Moving Average (EMA) decay of *α* = 0.999, where the EMA update rule follows:

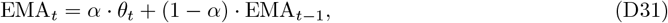

with *θ*_*t*_ representing the model parameters at iteration *t*. This EMA strategy stabilizes training by averaging recent parameter updates, reducing noise-induced fluctuations. For evaluation, the model is validated on a subset of antigen-antibody complexes, and the checkpoint achieving the highest TM-Score [**?**] is selected as the optimal model. The training task involves perturbing the antibody structure by introducing Gaussian noise (*σ* = 0.2 Å) to atomic coordinates, followed by reconstructing the perturbed structure. This process lasted for 5 days on 8 A100 GPUs, with gradient accumulation applied to simulate larger effective batch sizes when computational constraints limited the actual batch size.

Once the first-stage model has converged, we initialize the second stage using the parameters from the first-stage training. In this phase, the complexity is escalated by perturbing both the sequence and structure of the antibody’s Complementarity-Determining Regions (CDRs). Specifically, CDR sequences are mutated via profile-guided substitution (*e*.*g*., replacing residues with similar physicochemical properties), while structural perturbations are applied with higher magnitude (*σ* = 0.5 Å) to challenge the model’s ability to recover intricate interactions. The mixed training approach assigns probabilities of 4 : 2 : 2 : 2 for the model to design: (1) CDR H3 (the most structurally dynamic region), (2) CDR H1/H2, (3) all CDRs simultaneously, and (4) no sequence design (purely structural refinement). This distribution ensures the model learns hierarchical dependencies, starting from localized motifs (H3) to broader CDR interactions, while retaining the ability to handle cases where sequence information is absent. The second stage also lasted 5 days on 8 A100 GPUs, with additional data augmentation via random rotations and translations to further diversify the training manifold. Throughout both stages, self-conditioning [66] is implemented to improve training stability. This technique involves feeding the model with auxiliary information derived from the original (unperturbed) structure, such as backbone dihedral angles (*ϕ, ψ*) and hydrogen bond patterns. By conditioning on these low-level geometric features, the model learns to disentangle perturbation-specific variations from invariant structural principles, enhancing its capacity to generalize to unseen complexes. For instance, during reconstruction, the model explicitly aligns predicted side-chain conformations with the reference backbone, reducing overfitting to noisy perturbations.

After training the generative model, we distill the consistency model using the method proposed by [67]. This process involves training a lightweight student model to mimic the behavior of the teacher model (IgGM) across multiple noise levels, leveraging consistency regularization to enforce smoothness in predictions. Specifically, the student model is optimized to produce outputs that remain consistent under small input perturbations, ensuring robustness to sampling variability. The distillation loss combines mean squared error (MSE) between teacher and student outputs with KL divergence to align probability distributions, enabling efficient inference while preserving high-fidelity structural details. For specific implementation details, including hyperparameter tuning and ablation studies, please refer to [67].

### D.5 Evaluation Metrics

- Root Mean Square Deviation (RMSD): This metric obtained by calculating the mean of the squared differences between the coordinates of aligned CDR H3 backbone atoms and then taking the square root. The CDR H3 region is the most flexible, making it more challenging to predict.
- TM-Score (Template Modeling Score): This metric measures the structural similarity between the predicted antibody structure and a reference structure. A TM-Score of 1 indicates an exact match between the two structures, while a score closer to 0 indicates a poor match, a high TM-Score suggests that the predicted structure closely resembles the native structure.
- GDT-TS (Global Distance Test-Total Score): This metric provides an overall assessment of the model’s accuracy by comparing the predicted structure to the native structure based on the global distance test. It takes into account both the accuracy of the model’s predictions and the conformational similarity to the native structure. A higher GDT-TS score indicates a better match between the predicted structure and the native structure, suggesting a higher quality design.
- DockQ: This metric is specifically designed for evaluating the quality of protein-protein docking predictions. It assesses the interface complementarity and the conformational accuracy of the predicted complex structure. A high DockQ value indicates that the predicted interface is likely to be functional and stable, suggesting a well-designed multi-chain interface.

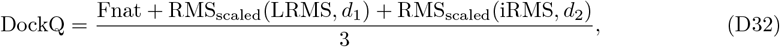

where RMS_scaled_ represents the scaled RMS deviation corresponding to either LRMS or iRMS, *d*_*i*_ is a scaling factor, *d*_1_ is used for LRMS, and *d*_2_ is used for iRMS. Fnat is defined as the fraction of native contacts retained in the predicted complex interface.
- SR (Success Rate): Indicates that the quality of the designed multi-chain interface positioning is within an acceptable range when DockQ is greater than 0.23. High SR indicates that more structure of antibodies is good.
- KD value (Dissociation Constant): It is used to describe the equilibrium dissociation constant of the binding between an antibody (Ab) and an antigen (Ag), reflecting the binding strength. Its calculation formula is *K*_*D*_ = [*Ab*][*Ag*]*/*[*Ab − Ag*]. The smaller the value (such as 10^*−*9^ M), the higher the affinity. Experimental methods include surface plasmon resonance (SPR), microplate detection (such as Octet), and isothermal titration calorimetry (ITC).
- EC50 (Half-maximal effective concentration): It refers to the antigen concentration required to achieve 50% of the maximum effect of the antibody. It is often evaluated through enzyme-linked immunosorbent assay (ELISA) or cell activity experiments (such as CBA, Luminex).

## E Wet Lab

### E.1 TNF*α* and IL-33

#### E.1.1 Expression and purification of human TNF*α* and IL-33

Recombinant human TFN*α* (hTNF*α*) was produced using Expi293 cells (ThermoFisher) via transient transfection with mammalian expression vector pcDNA3.1 (+), which contains the residues 77-233 and N-terminal signal peptide and 6xHis tag. Five days post-transfection, the supernatants were collected by centrifuging the culture at 3,900 rpm for 30 min and filtering through 0.45 *µ*m vacuum filter. Recombinant hTNF*α* was purified using Ni-NTA resin (Smart-Lifesciences) with buffer B (10 mM Na2HPO4, 10 mM NaH2PO4 [pH 7.4], 500 mM NaCl, and 250 mM imidazole). The eluted protein was subsequently concentrated and buffer-exchanged into PBS. Recombinant human IL-33 (residues 112-270) was expressed in Rosetta through transformation with the prokaryotic expression vector pET28a. The construct encoded an N-terminal 6×His tag fused to the mature IL-33 domain. Cells were cultured in SB medium supplemented with 100 *µ*g/mL kanamycin until reaching an OD_600_ 0.6, followed by induction with 1 mM IPTG at 16°C for 16 hr. Harvested cells were resuspended in PBS and lysed via high-pressure homogenization. The purification process using Ni-NTA resin (Smart-Lifesciences) with buffer B, subsequent concentration, and buffer exchange into PBS were performed as described for hTNF*α*. Protein purity was confirmed by SDS-PAGE analysis with concentration determination using the NanoDrop2000 spectrophotometer.

#### E.1.2 Expression and purification of sdAbs

The coding sequences of VHH3, I7 and their mutants (gene synthesis by GENEWIZ, Suzhou, China) were cloned into pComb3x expression vector containing an N-terminal OmpA secretion signal and C-terminal dual-affinity tags (6×His tag with FLAG tag tandem). Expression of these constructs was conducted in E. coli HB2151 at 30°C for 14 hours, induced by 1 mM isopropyl *β*-D-1-thiogalactopyranoside (IPTG). The bacterial cells were harvested and clarified lysates obtained by centrifugation at 8,000 ×g (4°C, 30 min) were subjected to immobilized metal affinity chromatography using HisTrap HP columns (Cytiva). The purification protocol included: (1) column equilibration with buffer A (10 mM Na2HPO4, 10 mM NaH2PO4, pH 7.4, 500 mM NaCl, and 20 mM imidazole), (2) 10-column volume wash with buffer A containing 25 mM imidazole, and (3) elution with 250 mM imidazole in buffer A. Purified proteins were dialyzed against PBS, snap-frozen in liquid nitrogen, and stored at -80°C.

#### E.1.3 ELISA

High-binding 96-well microplates (Corning #3690, half-area) were coated with recombinant human TNF*α* or IL-33 (100-200 ng/well in PBS, pH 7.4) overnight at 4°C. After three washes with PBS containing 0.05% Tween 20 (PBST), nonspecific binding sites were blocked with 5% (w/v) BSA in PBS (blocking buffer) for 1 hr at 37°C. Serial threefold dilutions of sdAbs were applied to antigen-coated wells and incubated for 90 min at 37°C. Following three PBST wash cycles, bound sdAbs were detected with horseradish peroxidase (HRP)-conjugated anti-FLAG M2 monoclonal antibody (1:5,000 dilution; Sigma-Aldrich #F3165) through 45 min incubation at 37°C. Colorimetric development was initiated by adding 2,2’-azino-bis (3-ethylbenzothiazoline-6-sulfonic acid) (ABTS) substrate solution (Thermo Scientific), the signal was read at 405 nm using a Microplate Spectrophotometer (Biotek). The EC50 values were extrapolated by fitting the data to a sigmoidal dose-response curve with a variable slope using GraphPad Prism version 8.0.

#### E.1.4 Biolayer interferometry (BLI) assay

The binding kinetics of single domain antibodies (sdAbs) to Protein A or interleukin-33 were evaluated by BLI using an Octet RED76e system (Sartorius AG). For VHH3 and its mutants binding to Protein A, threefold serial dilutions of antibodies were analyzed with ProA biosensors (Sartorius AG) through a three-phase protocol: (1) 120 s baseline equilibration in kinetics buffer (PBS containing 0.02% Tween 20), (2) 300 s association phase with diluted sdAbs, and (3) 300 s dissociation phase in kinetics buffer. Biosensors were regenerated between cycles using glycine-HCl (pH 1.5) for subsequent measurements. For IL-33 interaction analysis, AR2G biosensors (Sartorius AG) were first activated with 20 mM EDC/10 mM sulfo-NHS in water for 300 s, followed by covalent immobilization of 20 *µ*g/ml IL-33 in 10 mM sodium acetate (pH 5.0) until saturation After quenching with 1 M ethanolamine (pH 8.5) for 300 s, the functionalized sensors underwent baseline stabilization in kinetics buffer. Serial threefold dilutions of I7 and mutants were then analyzed through 300 s association and 300 s dissociation phases. Binding parameters were calculated using Data Analysis Software 11.1 (Sartorius AG) through global fitting with either 1:1 or 2:1 binding model.

### E.2 R1-32

#### E.2.1 Expression and purification of designed antibodies

The heavy chain (IgH) and light (IgL) chain genes of designed antibodies were amplified and cloned into the expression vector pCMV3. Expi293F cells (Thermo Fisher Scientific, A14527) were cultured in Expi293F culture medium (Thermo Fisher Scientific, A1435101) at 37 °C with 8% CO2 for protein expression. When the density of Expi293F cells reached 2.5 *×* 10^6^ cells/mL, the paired IgH and IgL plasmids were transiently co-transfected into the Expi293F cells at a ratio of 1:1 using the transfection reagent polyethylenimine (PEI). Five days post-transfection at 33 °C, supernatants were harvested by centrifugation and incubated with Protein A Resin (Genscript, China) at room temperature for 2 h to enable antibody binding. After the resin was washed with PBS (Gibco), pH 7.4, the antibodies bound to the resin were eluted using 0.1 M citric acid, pH 3.0, and neutralized immediately with an equal volume of 1 M Tris-HCl, pH 8.0, to maintain the pH within the neutral range. Subsequently, fractions containing antibodies were concentrated and buffer-exchanged into PBS using a 50 kDa MWCO Amicon Ultra filtration (Merck Millipore) at 4 °C. Purified antibodies were aliquoted, flash-frozen, and stored at *−*80 °C until needed.

#### E.2.2 Expression and purification of SARS-CoV-2 RBD

The coding sequence of SARS-CoV-2 RBD (residues 332–527), fused with an N-terminal *µ*-phosphatase signal peptide and a C-terminal hexahistidine tag, was cloned into the expression vector pcDNA3.1. Similar to antibody production, the RBD plasmid was transiently transfected into Expi293F cells using the transfection reagent polyethylenimine (PEI) when the cell density reached 2.5 *×* 10^6^ cells/mL. Five days post-transfection at 33 °C, the culture supernatant was collected by centrifugation and supplemented with 25 mM phosphate, pH 8.0, 300 mM NaCl, and 5 mM imidazole, and recirculated onto a HiTrap TALON crude column (Cytiva). Subsequently, the column was washed with buffer A (25 mM phosphate, pH 8.0, 300 mM NaCl, 5 mM imidazole), and the RBD protein was eluted with a 100 mL linear gradient to 100% buffer B (25 mM phosphate, pH 8.0, 300 mM NaCl, 500 mM imidazole). Fractions containing the target protein were pooled, concentrated with a 10 kDa MWCO Amicon Ultra filtration (Merck Millipore). Eventually, the RBD protein was further purified and buffer-exchanged into PBS using a Superdex 200 increase 10/300 GL column (Cytiva). All SARS-CoV-2 RBD variants were expressed and purified following the aforementioned protocol. Purified RBDs were aliquoted, flash frozen, and stored at *−*80 °C until needed.

#### E.2.3 Biolayer interferometry (BLI) binding assay

Binding kinetics and affinities of designed antibodies against the RBDs of wild-type SARS-CoV-2 and the variants were assessed by biolayer interferometry (BLI) on an Octet-RED76 (ForteBio). All experimental procedures were performed at 25 °C and all proteins were diluted to PBST buffer (PBS with 1 mg/mL BSA and 0.02% v/v Tween-20). Initially, antibodies (11 µg/mL) were immobilized onto Protein A biosensors. After a 60 s baseline step in PBST buffer, biosensors loaded with antibodies were exposed (300 s) to various RBDs (200 nM) to measure association. Subsequently, the biosensors were dipped into PBST buffer (600 s) to measure dissociation of RBDs from the biosensor surface. A blank reference was set for each reaction. Data was reference-subtracted and analyzed using the Octet Analysis Studio software v.12.2, fitting the data to 1:1 or 2:1 binding model to determine kinetic parameters. Results were plotted using GraphPad Prism 8.0.

### E.3 De novo design PD-L1 antibody

Programmed Death-Ligand 1 (PD-L1), a cell-surface receptor that interacts with PD-1, plays a key role in immune regulation and tumor immune escape. Here we introduce the relevant steps of wet experiments.

#### E.3.1 Antibody construction and HEK293 transient expression

The variable heavy (VH) and light (VL) chain fragments of scFv antibodies were amplified by PCR and purified using a membrane-binding buffer system. Purified fragments were ligated into the cloning vector with ligase at 50°C for 20 min, followed by transformation into competent cells via ice incubation (30 min) and heat shock (90 s). Colonies were expanded and sequenced to confirm scFv sequences. Validated heavy/light chain plasmid pairs were co-transfected into HEK293 cells using transfection reagent TF2. Transfected cells were cultured in serum-free CD medium supplemented with feed medium on days 1, 3, and 5 post-transfection, under conditions of 37°C, 5% *CO*_2_, and 175 rpm shaking.

#### E.3.2 Antibody purification and quality analysis

Cell supernatants were clarified by centrifugation (4,000g, 30 min) and filtered through 0.45 *µ*m membranes. Clarified samples were loaded onto protein A columns pre-equilibrated with AC Binding buffer. After washing (5–10 column volumes, CV), antibodies were eluted with AC Elution buffer, neutralized with Tris (pH 8.0), and dialyzed. Columns were regenerated with CIP buffer (*>*5 CV), rinsed to neutrality, and stored in 25% ethanol.

Antibody concentration was determined by measuring absorbance at 280 nm (baseline stability: ±0.015) and normalized using an IgG extinction coefficient (1.414). Purity was assessed via SDS-PAGE: samples (5 *µ*g) were denatured in 4× loading buffer, electrophoresed at 100 V (stacking gel) and 140 V (separation gel), and stained with Coomassie blue.

#### E.3.3 ELISA for antibody binding analysis

Microtiter plates were coated overnight at 4°C with antigen protein (0.1 or 1 *µ*g/mL, 100 *µ*L/well). After removing the coating solution, plates were blocked with 2% BSA (300 *µ*L/well) for 1 h at room temperature. Unbound blockers were removed by washing twice with 300 *µ*L/well of wash buffer, followed by thorough drying. Test antibodies (1 *µ*g/mL, 100 *µ*L/well) were added and incubated for 2 h at room temperature. Plates were washed three times with wash buffer, and horseradish peroxidase (HRP)-conjugated Goat Anti-Human IgG (H+L) secondary antibody (100 *µ*L/well at optimal dilution) was added for 1 h. After repeating the wash step, chromogenic substrates A and B were mixed (1:1 ratio) and added to each well (200 *µ*L/well). The reaction proceeded for 20 min at room temperature in the dark before termination with 50 *µ*L/well stop solution. Absorbance at 450 nm (OD450) was measured immediately.

## F Ablation Study

We conducted ablation experiments on parts of the model and evaluated it on the test set, as shown in Table B5. The removal of the two-stage training resulted in a significant decline in all metrics, particularly with the SR dropping to 0, indicating that the two-stage training is crucial for model performance. The absence of epitope information led to a marked deterioration in the DockQ, iRMS, and LRMS metrics, highlighting the importance of epitope information on the docking quality and bias of the model. After removing PPSM, there was a slight decrease in AAR and DockQ, while iRMS and LRMS increased, suggesting that PPSM has a certain impact on the bias of the interface and ligand. The removal of mixed training also resulted in a slight decline in AAR and DockQ, with increases in iRMS and LRMS, indicating that mixed training has a notable effect on the overall performance of the model. Overall, the various components and training strategies of the IgGM model significantly influence its performance, with the two-stage training and epitope information being particularly critical for the docking quality and success rate of the model.

## G Additional Results

To explore the diversity of antibodies designed by IgGM and dyMEAN, we conducted a statistical analysis focusing on antibody designs in which the CDR H3 region had an amino acid sequence length of 14 (Fig. A1 b), the most abundant lengths [39]. The distribution of lengths is presented in Fig. A1 a. The plot shows that the distribution of experimental antibody sequences is fairly even for antibodies with this specific length. The CDR H3 region is the area with the highest relative abundance of each amino acid in the entire antibody [39]. Compared to native antibodies, both IgGM and dyMEAN designed antibodies tend to generate a higher frequency of the amino acid tyrosine (Y). This preference can be attributed to the biological significance of tyrosine, which constitutes approximately 18.6% of the residues in the CDRH3 region according to the OAS database. Tyrosine is particularly critical in the antigen-binding site (PCS), where it is the most abundant amino acid residue, accounting for 19% [90]. Tyrosine provides multiple contact points that facilitate the positioning of the epitope-containing surface (ECS) of the antigen on the PCS. However, the sequences generated by dyMEAN are relatively homogeneous, while those produced by IgGM exhibit greater diversity, which can be attributed to the variability inherent in the diffusion model. We further analyzed the distribution of amino acid sequences in the CDR3 region of heavy chains within the test set and compared the sequence distributions associated with dyMEAN and IgGM. dyMEAN tends to generate a high frequency of the amino acids glycine and tyrosine, with their combined distribution approaching nearly half of the total generated sequences, whereas in reality, they account for only about one-quarter. For some less common amino acids, such as glutamine and tryptophan, dyMEAN did not generate any instances. In contrast, the sequence distribution designed by IgGM aligns more closely with the distribution observed in the test set, showing a better fit for the frequencies of Glycine and Tyrosine, and maintaining a distribution that closely resembles the true distribution for each amino acid type. This also aligns more closely with the actual distribution of antibody sequences, as illustrated in Fig. A1 c. We present additional examples of the *de novo* designs generated by our IgGM in Fig. A4.

